# Reduction of nucleolar NOC1 accumulates pre-rRNAs and induces Xrp1 affecting growth and resulting in cell competition in *Drosophila*

**DOI:** 10.1101/2021.07.06.451100

**Authors:** Francesca Destefanis, Valeria Manara, Stefania Santarelli, Sheri Zola, Marco Brambilla, Giacomo Viola, Paola Maragno, Ilaria Signoria, Gabriella Viero, Maria Enrica Pasini, Marianna Penzo, Paola Bellosta

**Author notes:** equally contributed (Francesca Destefanis, Valeria Manara, Stefania Santarelli).

## Abstract

*NOC1* is a nucleolar protein necessary in yeast for both transport and maturation of ribosomal subunits. Here we show that in *Drosophila* NOC1 is essential for the correct animal development, and that its ubiquitous downregulation results in small larvae with reduced polysome abundance and decreased protein synthesis. NOC1 expression in multiple organs, such as the prothoracic gland and the fat body, is necessary for proper organ functioning. Reduction of NOC1 in clones from the imaginal discs results in small clones with cells that die by apoptosis, an event that is partially rescued using a *M/+* background, suggesting that reduction of NOC1 causes the cells to acquire a loser state. This event was supported also by an increase in the transcription of Xrp1 and by activation of the pro-apoptotic eiger-JNK pathway, resulting in the upregulation of DILP8 as an indication of cellular damage. Here, we show that *Drosophila* NOC1 is important in the control of pre-rRNAs maturation and essential step in the regulation of ribosome biogenesis and its downregulation results in defects in growth and in cell competition, highlighting its novel function in this field.

**summary statement:** NOC1 is a nucleolar protein necessary for protein synthesis and ribosomal assembling. Its modulation induces cell competition and affects animal growth.

## Introduction

NOC1, 2 and 3 are members of a large family of conserved nucleolar proteins that play a critical role in the control of ribosome biogenesis in yeast and plants (Edskes et al., 1998; Li et al., 2009). Studies in *S. cerevisiae* reveal that NOCs proteins are required for the maturation and processing of the rRNAs (Khoshnevis et al., 2019) and for transport of the pre-ribosomal 60S subunit in the cytoplasm through the formation of NOC1/NOC2 and NOC2/NOC3 heterodimers (Hierlmeier et al., 2013; Milkereit et al., 2001). NOC1-3 function is unique and essential, as mutation in each gene affects growth and viability in both *S. cerevisiae* and in *Arabidopsis* (Edskes et al., 1998; Li et al., 2009; Milkereit et al., 2001).

In *Drosophila,* efficient ribosome biogenesis is necessary during larval development, when increase in cell mass and animal size is highly dependent on protein synthesis (Texada et al., 2020). Mutations in genes that regulate this process, like those encoding for *Minute* ribosomal proteins (Marygold et al., 2007; Saeboe-Larssen et al., 1998) or for *Nop60b/Dyskerin* (Tortoriello et al., 2010) and *Nopp140* (Baral et al., 2020), components of the nucleolus, present common defects that include a delay in development and reduced body size. Similar phenotypes have also been described for mutations in genes that control rRNA synthesis, such as the RNA-Pol-I associated chromatin regulator *PWP1* (Liu et al., 2017) or the *Rpl-135* subunit of the Pol-I complex (Grewal et al., 2005), and for *diminutive* (*dm*), the gene encoding for MYC (Johnston et al., 1999), a master regulator of ribosome biogenesis both in *Drosophila* and in vertebrates (Barna et al., 2008; Destefanis et al., 2020; Grewal et al., 2005; van Riggelen et al., 2010).

Larval growth is also regulated by *Drosophila* insulin-like peptides DILPs (DILP2, 3 and 5) released from the Insulin Producing Cells (IPCs) in response to nutrients (Geminard et al., 2009; Koyama et al., 2020; Maniere et al., 2020). This process is developmentally coordinated by the growth hormone ecdysone, secreted by the ring gland (Nijhout et al., 2014), and indirectly by DILP8, a peptide member of the Insulin/Relaxin family, secreted by cells from the peripheral organs in response to tissue damage (Garelli et al., 2015; Vallejo et al., 2015). The release of DILP8 blocks ecdysone synthesis and delays development to ensure proper regeneration of the damaged organ with developmental timing (Boulan and Leopold, 2021). In cells of the imaginal discs, DILP8 upregulation has been associated with cell damage induced by the activation of eiger/JNK pathway (Sanchez et al., 2019), and more recently with the transcriptional upregulation of the Xrp1*-*RpS12 axis (Boulan and Leopold, 2021) that links signals of inter-organ coordination with proteotoxic stress. Indeed, reduced protein synthesis activates a stress response which involves the activation of Xrp1, a pro-apoptotic CCAAT-Enhancer-Binding Protein (C/EBP) that eliminates the unfitted cells by cell competition (Baillon et al., 2018; Brown et al., 2021). Proteotoxic stress is induced by mutations in ribosomal proteins, such as RpS3 (Akai et al., 2021; Baumgartner et al., 2021) and RpS12 (Ji et al., 2019; Lee et al., 2018), or by the activation of the JNK/STAT signaling pathway (Kucinski et al., 2017), revealing the complexity of this network signaling.

In this study we characterized the function of *Drosophila* nucleolar NOC1, NOC2 and NOC3 *in vivo* and showed that their activity is necessary for proper animal growth. We demonstrated that NOC1 controls polysome abundance and its ubiquitous reduction blocks rRNA maturation resulting in reduced protein synthesis. In line with these results, lowering NOC1 levels in the whole animal results in small larvae that die early during development, which agrees with the observation that its reduction in different organs causes specific functional impairments. In cells of the wing imaginal disc, NOC1 downregulation induces apoptosis that is rescued in a *Minute/+* background and by P35 expression. The increase of Xrp1, eiger and JNK pathways reveals the activation of mechanisms that link NOC1 reduction to the induction of proteotoxic stress leading to cell death and cell competition.

## Results

### *Drosophila* NOC1 localizes in the nucleolus and is necessary for animal growth

*NOCs* (Nucleolar complex associated) are members of a protein family characterized by the presence of a NOC domain, not conserved in all proteins, and necessary for their heterodimerization (Milkereit et al., 2001). In *Drosophila,* the orthologues of yeast *Noc1*, *Noc2* and *Noc3* are annotated as *CG7839, CG9246* and *CG1234,* and are hereinafter called *NOC1, NOC2 and NOC3.* These genes, present also in humans, have a grade of conservation that varies from 32% to 35% of homology within their amino acid sequences (Figure 1A, Supplementary Figure S1). Interestingly, a network analysis using the STRING database on predicted protein-protein interactions for NOC1/CG7839 uncovers that all three NOC proteins form a hub with other nucleolar proteins with a distinct role in ribosome biogenesis, suggesting that NOCs may function in concert to ensure proper nucleolar activity (Supplementary Figure S2).

**Figure 1:**
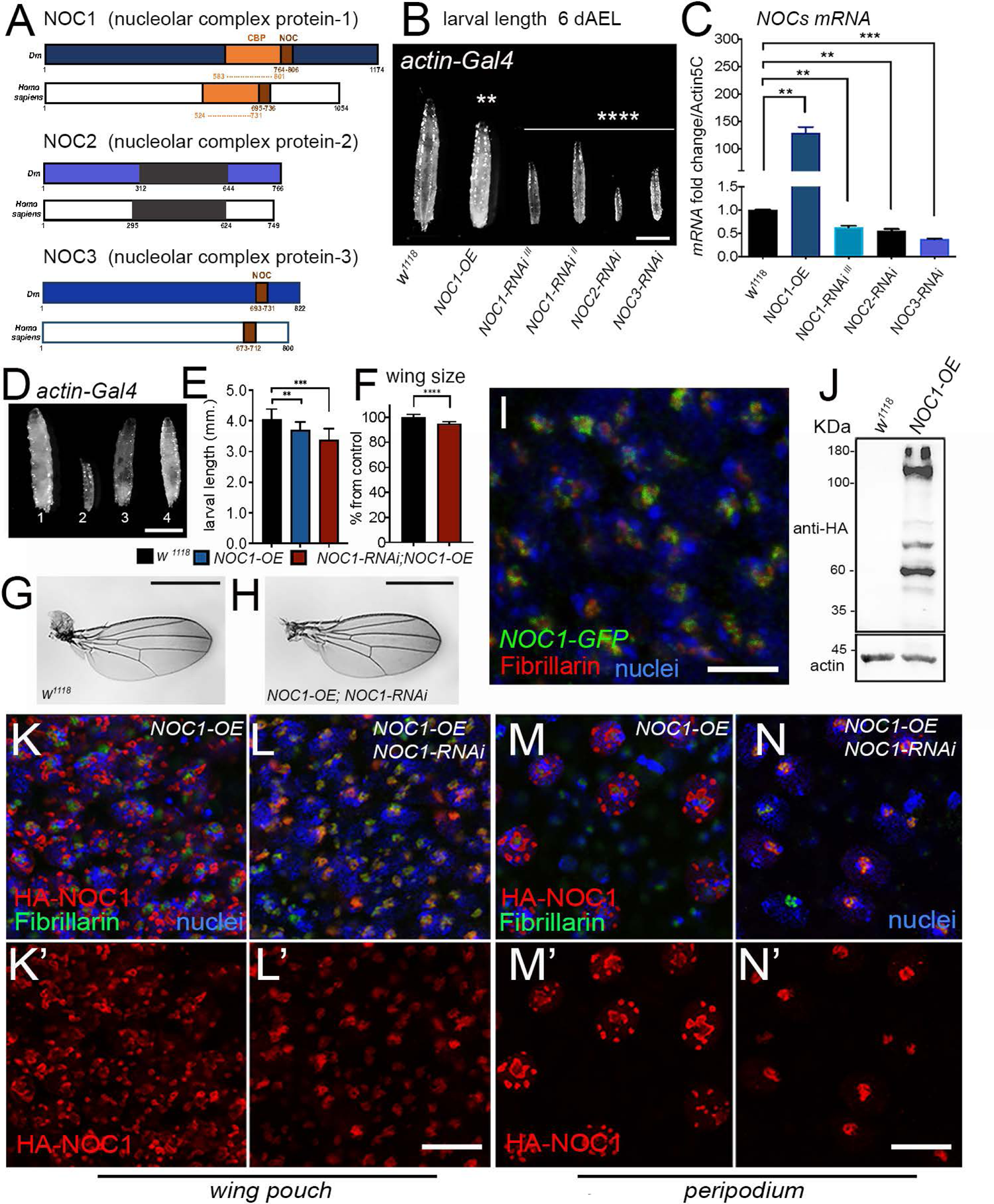
NOC1 expression in the nucleolus must be at proper levels to allow animal survival. (A) Schematic representation of *Drosophila* NOC1, NOC2 and NOC3 proteins and their human homologues. NOC1 proteins contain a CBP domain (CCAAT binding domain), in orange, that shares 32% identity between sequences. In brown is represented the conserved NOC domain of 45 amino acids, present only in NOC1 and NOC3, that share 48% and 38% sequence identity between Dm and human proteins respectively. NOC2 protein shares an overall 36% identity between Dm and human proteins and in black is represented the region of highest homology (48%). (B) Photos of third instar larvae expressing the indicated transgenes under the *actin* driver taken at 120 hours after egg laying (AEL); the scale bar represents 1 mm. (C) qRT-PCR showing the amount of the indicated *NOCs-mRNAs* from larvae at 120 hours AEL; *actin5C* was used as control. (D) Photographs of larvae at 120 hours AEL: (1) control *w^1118^*, (2) *NOC1-RNAi*, (3) *NOC1* overexpression (OE), (4) *NOC1-RNAi; NOC1-OE* using the *actin-Gal4* driver. (E) Larval length at 120 hours AEL. (F) Quantification of the wing’s area/size in animals of the indicated genotype; the number is expressed as % from the control *actin-w^1118^*, the error bars indicate the standard deviations. At least 10 animals were used for each genotype, experiment was repeated twice. (G) Photos representing wings from females of the indicated genotype; the scale bars in D, G and H represents 1 mm. (I) Confocal image of cells from the imaginal disc showing the expression the *NOC1-GFP* visualized using anti-GFP antibodies in green and anti-fibrillarin in red; nuclei are visualized with Hoechst; scale bar represents 5 μm. (J) Western blot analysis from lysates of larvae expressing *HA-NOC1* under the *actin* promoter showing a band of 132 KDa of the expected size for NOC1 is visualized by anti-HA antibody with few other bands at lower molecular weight; actin is used as control loading. (K-N) Confocal pictures of cells from the imaginal discs (K-L) or from the peripodial epithelium (M-N) expressing *HA-NOC1* (K-M) or *HA-NOC1; NOC1-RNAi* (L-N) using the *engrailed-*promoter. NOC1 expression was visualized using anti-HA antibodies in red and anti-fibrillarin in green. Statistical analysis in B, C and E was calculated using one-way ANOVA with Tukey multi-comparisons test from at least 10 animals ** = *p* < 0.01, *** = *p* < 0.001 and **** = *p* < 0.0001, and the error bars indicate the standard deviations. Statistical analysis in F was calculated using Student’s *t*-test from at least 10 animals ** = *p* < 0.01, *** = *p* < 0.001 and **** = *p* < 0.0001, and the error bars indicate the standard deviations.

Our results show that ubiquitous reduction of NOC1, 2 or 3 in *Drosophila*, using RNAi interference in combination with the *actin* promoter (Supplementary Figure S3), resulted in small larvae that die between first and second instar. On the contrary, overexpression of NOC1 led to larvae that reach pupariation at a smaller size than control and failed to maturate into adult animals (Figure 1B, C and Table 1). Next, we explored the hypothesis that ubiquitous expression of NOC1 could rescue the larval defects induced by NOCs-RNAi. This analysis showed that co-expression of NOC1 compensates for NOC1 reduction allowing larvae to reach metamorphosis (Figure 1D, E) and to mature into small but viable adults (Figure 1F, G-H). On the contrary, co-expression of NOC1 failed to rescue the lethality of NOC2-RNAi and NOC3-RNAi animals, indicating that its function does not complement for the NOC2 or NOC3 activities (not shown). To analyze whether NOC1 localizes in the nucleolus we took advantage of the line CG7839-GFP.FPTB generated in modENCODE (Model Organism ENCyclopedia Of DNA regulatory Elements), in which GFP-CG7839 is expressed under the control of its 5’-end regulatory sequences (Kudron et al., 2018). This analysis in cells of the wing imaginal disc showed NOC1-GFP expression primarily in the nucleolus, colocalizing with fibrillarin (Figure 1I). The same analysis was performed in cells in the salivary glands, confirming the colocalization between fibrillarin and NOC1-GFP (Supplementary Figure S4). Since no commercial antibodies are available for the endogenous *Drosophila* protein, we determined NOC1 molecular weight by expressing an HA-tagged form. This data showed that HA-NOC1 protein is expressed as a 132 KDa protein in lysates from 3rd instar larvae. In addition, we observed multiple other bands at lower molecular weights (Figure 1J), suggesting that HA-NOC1 may undergo to unusual proteolytic processes at this stage of development linked to the lethality observed during larvae and pupae transition (Table1). Overexpression of HA-NOC1 using the *engrailed* promoter resulted in a nucleolar staining that colocalized with fibrillarin, detected in the small cells of the columnar epithelium (Figure 1K-K’) and better defined in the large cells composing the peripodium (Figure 1M-M’). In addition, we also observed the formation of large nuclear granules outside the nucleolar zone in both cells, better visible in the nuclei of the peripodium’s cells (Figure 1M-M’). These large structures and the abnormal nucleolar morphology were rescued when NOC1-RNAi was expressed with HA-NOC1 (Figure 1L-N).

**TABLE 1.**
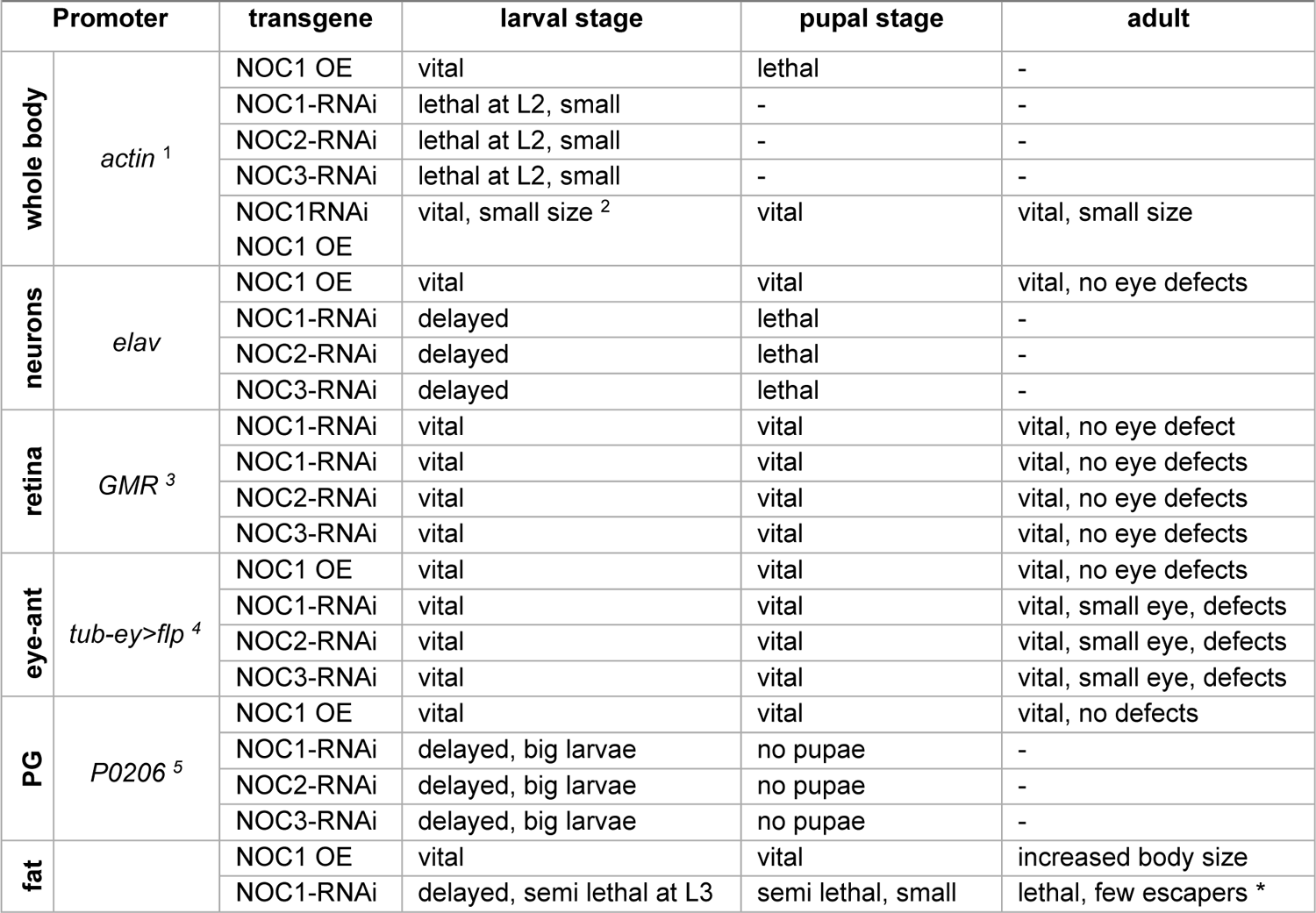
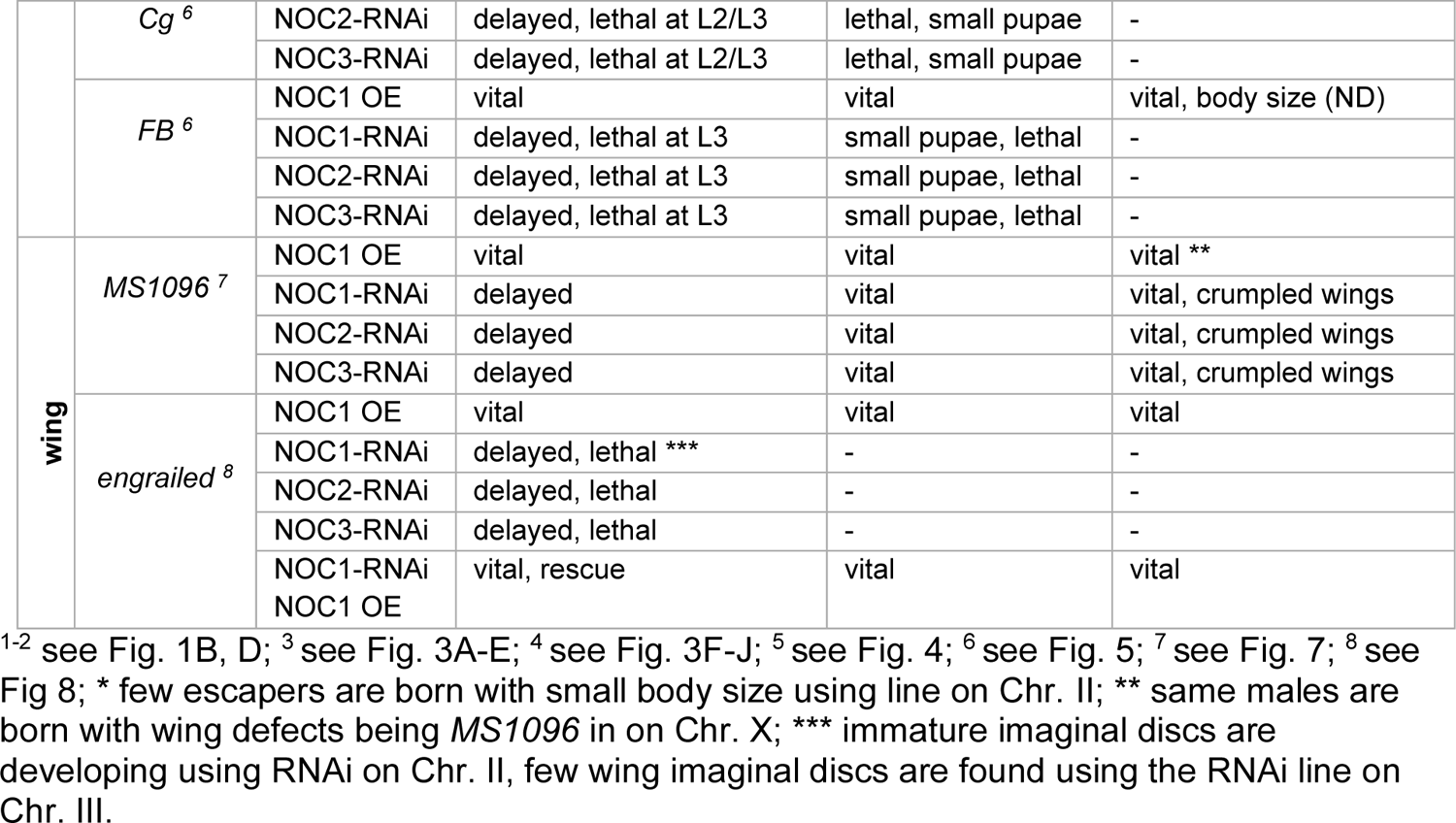
Characterization of NOCs expression *in vivo* using different promoters

### NOC1 is central for rRNA processing, ribosome maturation and functional protein synthesis

To investigate the role of *Drosophila*’s NOCs in ribosome biogenesis, we first analyzed the impact of NOC1 on ribosome maturation and protein synthesis. Polysome profiling in whole larvae showed that overexpression of NOC1 significantly increases the abundance of the 80S and polysomes peaks are increased compared to the WT (Figure 2B). On the contrary, NOC1 reduction resulted in a dramatic decrease in ribosomal subunits and polysomes abundance (Figure 2C) with a robust reduction of the 80S, and the relative increase of the 40S and 60S subunits, suggesting a defect also in proper ribosome recruitment on polysomes (Figure 2D, E). In yeast, the NOC1/NOC2 interaction is necessary to regulate the activity of Rpr5, a fundamental assembly factor that blocks the premature cleavage of the internal transcribed spacers (ITS) sequences during maturation of the rRNA precursors. This process allows the processing of the 60S and the production of the two subunits in stoichiometric amount (Khoshnevis et al., 2019). To better asses if in *Drosophila* NOC1 controls this process, we evaluated the levels of ITS1 and ITS2 and of the relative RNAs maturation by qRT-PCR. This analysis showed that reduction of NOC1 induces the accumulation of the intermediate ITS1 and ITS2 immature forms of rRNAs with the consequent reduction of the 18S and 28S rRNAs (Figure 2F). NOC1 overexpression instead reduces the level of ITS1 but not of ITS2, and significatively increases the amount of 18S and 28S rRNAs (Figure 2F), suggesting that NOC1 is part of the mechanism that controls rRNAs synthesis and processing in *Drosophila*. To evaluate whether this defect reflets in changes in global protein synthesis levels, we performed a SUnSET (Surface Sensing of Translation) assay (Deliu et al., 2017) in larval tissues. These experiments showed that translation of labeled puromycin-peptides is robustly diminished in *NOC1-RNAi* compared to control animals (Figure 2G-H). On the contrary, overexpression of NOC1 did not significantly impair the translation (not shown).

**Figure 2:**
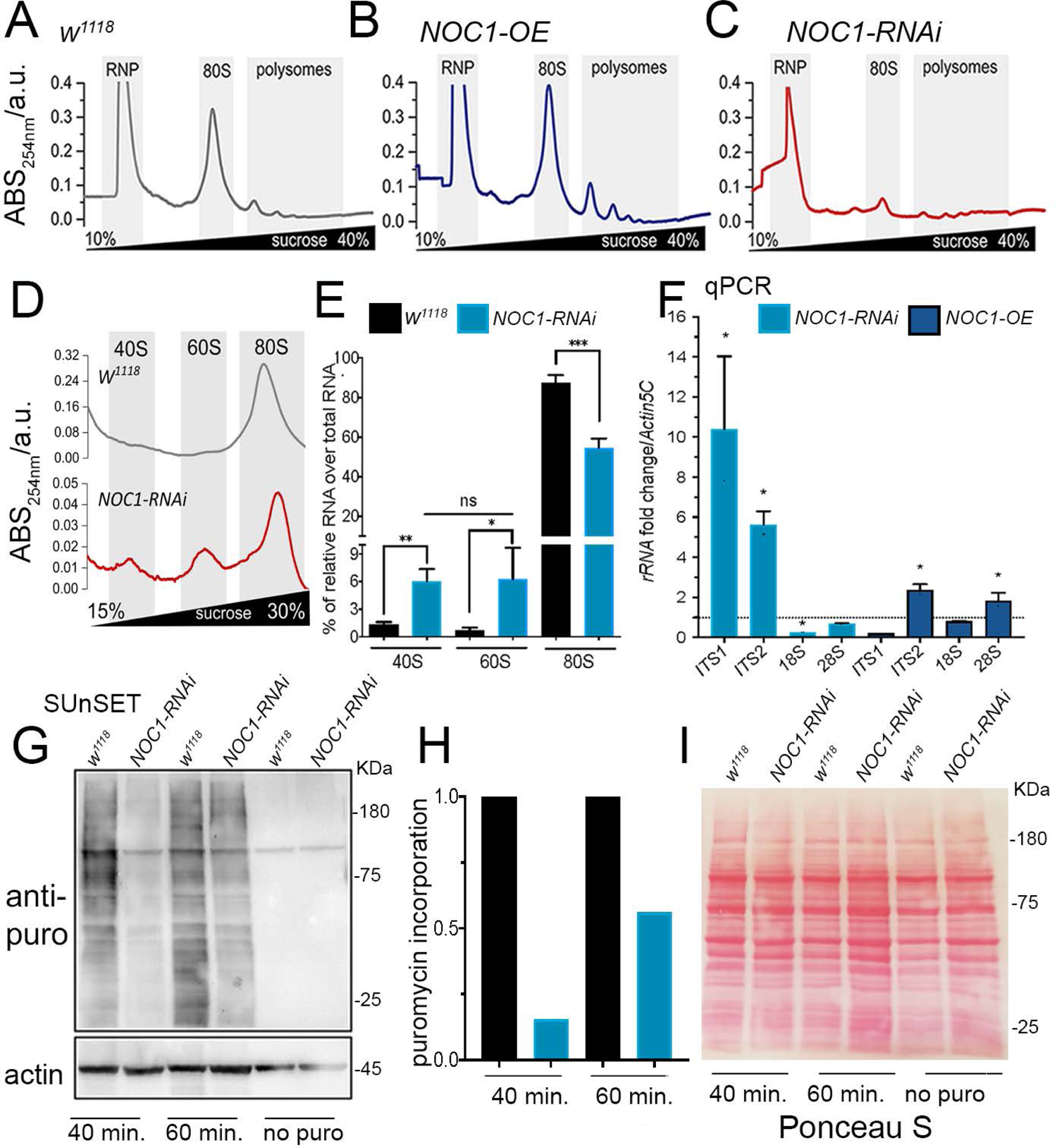
NOC1 regulates rRNA processing and ribosomal assembling, affecting protein synthesis. (A-C) Representative sucrose density gradient profiles of ribosome from control larvae (A) or animal over-expressing *NOC1* (B) or *NOC1-RNAi* (C). (D) Higher resolution of image A and C highlighting the area of the 40, 60 and 80S ribosomal subunits, note that the graphs use different scales. (E) Analysis of the % of 40, 60 and 80S ribosomal subunits, relative to each genotype, calculated over the total area including the polysome. (F) qRT-PCR showing the fold of induction over control *w^1118^* of pre-rRNAs analyzed using the ITSs (internal transcribe sequences) and of mature ribosomal rRNAs; data are expressed over *actin5C* used as control. (G) SUnSET western blot analysis of lysates from larvae treated with puromycin for the indicated time. The blot shows the relative changes in protein synthesis using anti-puromycin antibodies in control *w^1118^* or in larvae ubiquitously expressing *NOC1-RNAi* under the *actin*-promoter. Actin was used as control loading. (H) Quantification of the change in puromycin incorporation from G and normalized over actin (Deliu et al., 2017). (I) Ponceau S staining showing total protein levels in G. Statistical analysis in E and F was calculated using one-way ANOVA with Tukey multi-comparisons test from at least three independent experiments ** = *p* < 0.01 and *** = *p* < 0.001; the error bars indicate the standard deviations.

### Reduction of NOC1, NOC2 and NOC3 levels during development costrains growth in the eye but does not affect morphology in differentiated ommatidia

Next, we analyzed the function of NOC1 in tissues with different proliferative characteristics. Using the *GMR* promoter (Hay et al., 1994), we modulated NOC1 expression at mid-third instar stage in the differentiated cells of the retina, and using the *tubulin* promoter in combination with *eyeless-*flippase, we restrained the expression of NOC1 to the proliferative cells precursors of the eye and antenna discs (Bellosta et al., 2005). These experiments showed that downregulation of NOCs or overexpression of NOC1 in the differentiated cells under the *GMR* promoter did not affect the eye morphology (Figure 3A-E). On the contrary, using the *tubulin* promoter, downregulation of NOC1, 2 and 3 resulted in small eyes with fewer and disorganized ommatidia (Figure 3H-J), whereas overexpression of NOC1 did not change the eye morphology (Figure 3G). Moreover, the eye defects induced by NOC1-RNAi can be rescued by co-expression with the inhibitor of caspase P35 (Figure 3L-M), suggesting that cell death is responsible for these defects.

**Figure 3:**
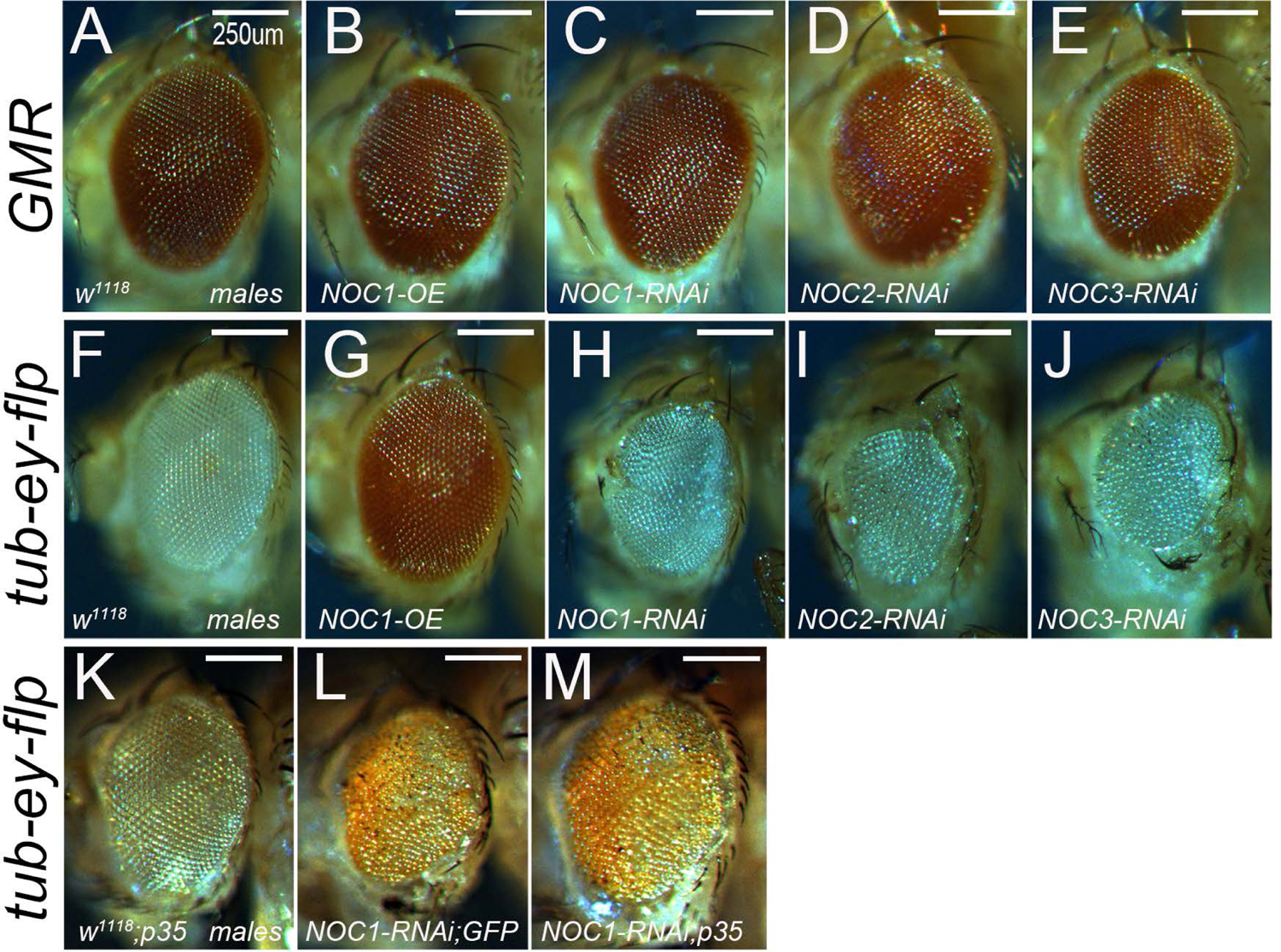
Reduction of NOC1, NOC2 and NOC3 affects the development of the eye and antenna and induces apoptosis but does not affect the differentiated ommatidia. Photographs of *Drosophila* compound eyes (lateral view) expressing the indicated transgenes using the *GMR-Gal4* promoter (A-E) or the *tubulin-GAl4* promoter in combination with *eyless-flippase* to constrains Gal4 expression in the proliferative cells of the eye and antenna (F-J) (Bellosta et al., 2005). (K-M) Photographs of compound eyes expressing the caspase inhibitor *P35* alone (K) or together with *NOC1-RNAi* (M) that rescues the eye defect showed in L. The scale bars represent 250 μm. Photographs were acquired from male’s eyes and similar data were obtained using females (not shown).

To further characterize the role of NOCs *in vivo* we evaluated the impact of its downregulation or overexpression in organs important for the physiological growth of the animal.

### Reduction of NOC1 in the prothoracic gland affects ecdysone levels, inhibiting pupariation

Downregulation of NOCs in the prothoracic gland (PG) using the *P0206-Gal4* promoter delayed larval development. These animals did not pupariate after 5 days AEL but continued to grow and feed for about 20 days before dying (Figure 4F). Macroscopic analysis of the PG revealed that at 5 days AEL its size in *NOC1-RNAi* larvae was similar to that of controls, while in *NOC1-RNAi* animals at 12 days AEL was smaller than the one in animals at 5 days AEL (Figure 4A-C). Cells in the PG are specialized for ecdysone synthesis, thus we determined its production by measuring the level of its target *Ecdysone-induced protein 74b (E74b)* from whole larval tissues (Valenza et al., 2018). Our data showed that the level of *E74b mRNA* was reduced at 5 days AEL in *NOC1-RNAi* animals compared to controls and was further lowered at 12 days AEL (Figure 4D). *NOC1-RNAi* expressing larvae keep wandering and increased their larval size (Figure 4E) while the cells of the fat body increased volume and fats (not shown). On the contrary, NOC1 overexpression did not lead to detectable changes in *E74b mRNA* levels or in body size (not shown).

**Figure 4:**
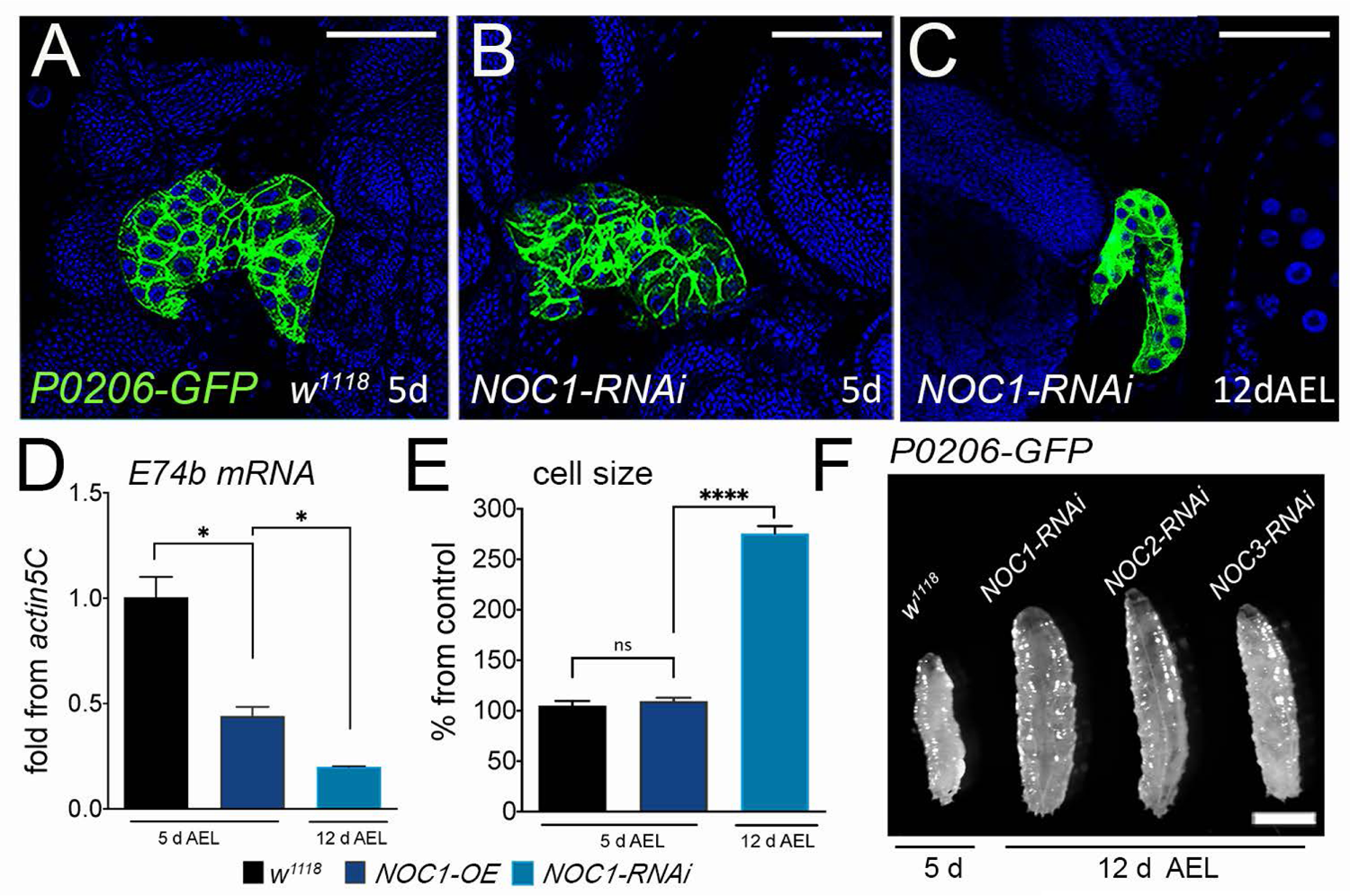
NOC1 downregulation in the prothoracic gland reduces ecdysone production and delays development. (A-C) Confocal images of the ring gland marked with GFP, using the *P0206-GFP* driver line, from control *w^1118^* (A) and from animals with reduced *NOC1* at 5 days AEL (B) and 12 days AEL (C). Nuclei are stained with Hoechst; the scale bar represents 50 μm. (D) qRT-PCR showing the level of *E74b* mRNA, target of ecdysone, from whole larvae at the indicated time of development. (E) Analysis of cell size in the fat body from control *w^1118^* or *NOC1-RNAi* larvae at 5 days and at 12 days AEL. (F) Photographs of whole animals with reduced NOCs expression in the prothoracic gland. Picture represents *w^1118^* larvae at 5 days AEL and *NOC1, 2 and 3-RNAi* at 12 days AEL. The scale bar represents 1 mm. Statistical analysis in D and E was calculated using one-way ANOVA with Tukey multi-comparisons test from at least two independent experiments, more than 10 animals were used in each experiment * = *p* < 0.05 and **** = *p* < 0.0001; the error bars indicate the standard deviations.

### NOC1 downregulation in the fat body reduces cell size and lipid storage causing dyslipidemia

*NOC1, 2,* or *3* downregulation in clones of the fat body affected cell morphology and significantly reduced cell size (Figure 5C-E, F). On the contrary, NOC1 overexpression did not change cell-morphology but resulted in a small decrease in the total area of the clones (Figure 5B, F). To investigate the impact of NOC1 expression in the whole organ, we used the *Cg (Collagene4a1)* promoter, that is active in the fat body and in the hemocytes, and the *FB* promoter for specific expression in fat cells (Schmid et al., 2014). Reduction of NOCs, with both promoters, results in larvae with a small body size that delayed their development and died at either late third instar or pupal stages (Table 1). We noticed that the developmental delay was less pronounced using the line *NOC1-RNAi ^II^* where a small percentage of animals (<10%) hatched as small adults (Figure 5H, Table 1). A possible explanation for this difference may be the lower efficiency of the *NOC1-RNAi ^II^* line in reducing *NOC1-mRNA* compared to the line on the Chromosome III (Supplementary Figure S5). Overexpression of NOC1 induced an increase in larval size that was significant at 96AEL, resulting to adults that hatched slightly bigger than control as shown by an analysis of their wing size (Figure 5H).

**Figure 5:**
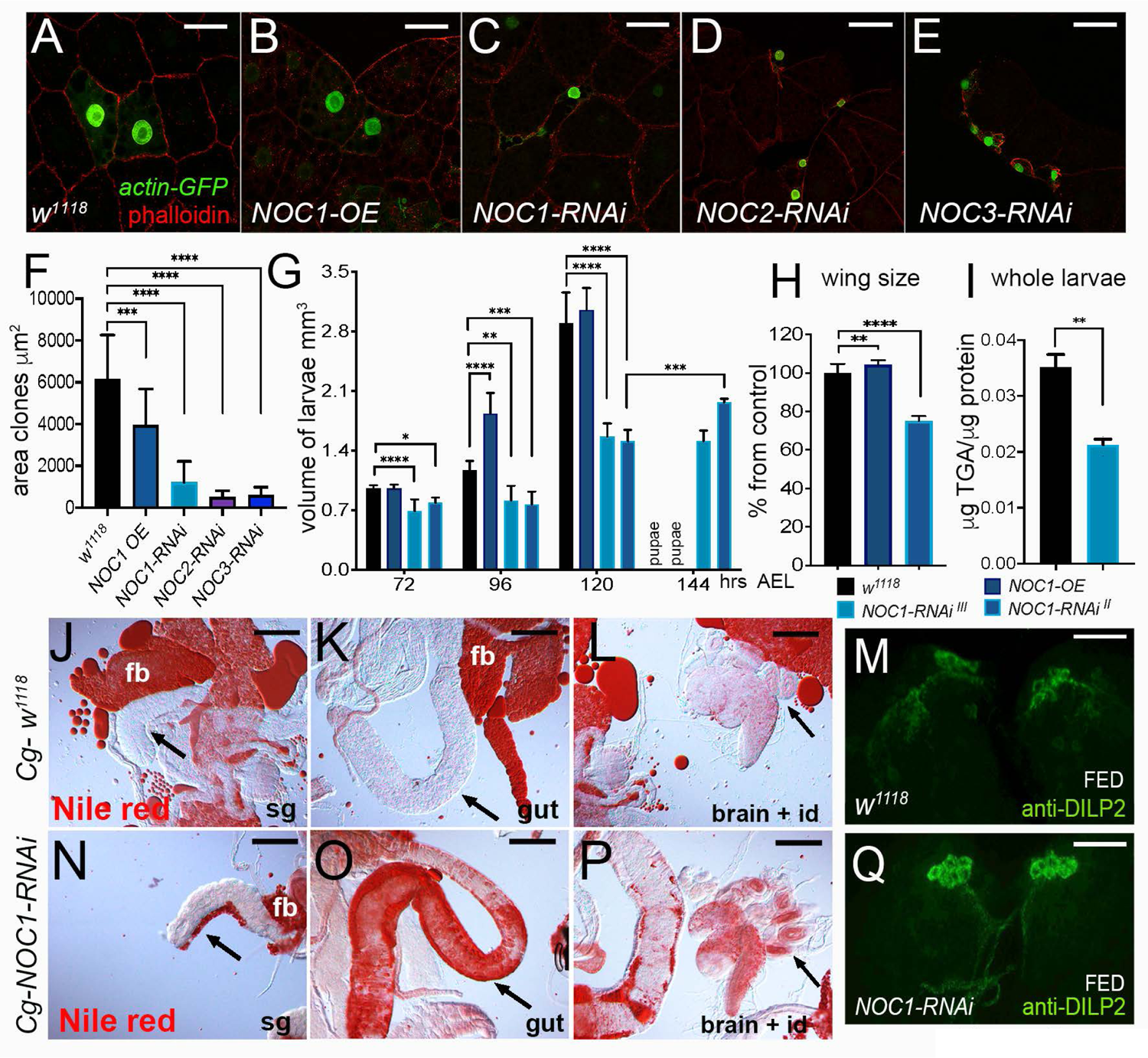
NOC1 downregulation in the fat body reduces cell size and induces dyslipidemia. (A-E) Confocal images of *actin-*flip-out clones in the fat body co-expressing nuclear GFP together with the indicated transgenes. Phalloidin-Texas Red was used to mark cell membranes; the scale bars represent 50 μm. (F) Quantification of the area of the clones in the fat body. (G) Analysis of larval volume measured at the indicated time of development until pupariation in animals in which the NOCs transgenes were expressed using the *Cg* promoter. (H) Analysis of the wing size from four days old females of the indicated genotypes, data are expressed as % from control *w^1118^*. (I) Quantification of triglycerides (TGAs) in whole larvae at 120 hours AEL, data are expressed as μg of TGAs/μg of proteins. (J-L, N-P) Photographs of larval organs stained with Nile red to visualized lipids from control *w^1118^* (J-N) and NOC1-RNAi animals (N-P) at third instar. (J-N) Reduction of *NOC1-RNAi* affects the size of the fat body (fb) particularly visible near the salivary gland (sg indicated by the arrow); the scale bar represents 100 μm. The impairment to accumulate nutrients in the fat body in *NOC1-RNAi* animals induces the storage of fats in other organs, visible in the gut indicated by the arrow in K and O, and in the brain and imaginal discs (id), indicated by the arrow in L and P. (M, Q) Confocal images of third instar larval brains showing DILP2 immunostaining in the Insulin Producing Cells (IPCs) from control *w^1118^* (M) and *NOC1-RNAi* (Q) animals in feeding conditions; the scale bar represents 50 μm. Data in G are representative of one of three experiments using ten or more animals for each genotype; data in F, G and H are calculated using one-way ANOVA with Tukey multi-comparisons test from at least two independent experiments; data in I were calculated using Student’s *t* test from two independent experiments; * = *p* < 0.05, ** = *p* < 0.01, *** = *p* < 0.001and **** = *p* < 0.0001, the error bars indicate the standard deviations.

The fat body stores lipids necessary for the animal to develop and to bypass metamorphosis (Edgar, 2006). Analysis of the triglycerides content (TGA) showed that *NOC1-RNAi ^III^* larvae have less lipids in their fat cells compared to wild type animals (Figure 5I, J-P). Morphologically, this can be observed in the fat body near to the salivary glands (sg), where it is almost absent in *NOC1-RNAi* animals (Figure 5N arrow). As a response for the reduced lipid storing capability, these animals suffered from dyslipidemia, an inter-organ process induced by the failure of fat cells to store lipids. This process is activated when reduced or impaired lipid metabolism produces molecules that non-autonomously induce the accumulation of lipids in other organs (Palm et al., 2012). Indeed, we observed that *NOC1-RNAi* animals accumulated high level of lipids in the gut (Figure 5K and O), in the brain and in the imaginal discs (Figure 5L and P).

The fat body also remotely controls the release of *Drosophila* insulin-like peptides (DILP2, 3 and 5) from the Insulin Producing Cells (IPCs), that are secreted in the hemolymph in response to nutrients or reteined during starvation (Geminard et al., 2009). Analysis of DILP2 expression in the IPCs showed that, even in adequate nutrients conditions (FED), DILP2 is retained in the IPCs of animals with reduced NOC1 in the fat body (Figure 5Q-M), suggesting that these animals lost their ability to remotely control the release of DILPs, thus mimicking starvation, a condition in which DILPs would ordinarily be retained.

### NOC1 reduction in cells of the wing imaginal disc induces apoptosis partially rescued in a Minute/+ background

To assess the impact of NOCs in the growth of epithelial cells, we generated flip-out clones where NOCs level was either reduced or overexpressed. Analysis of the clones performed at 72-90 hrs AEL showed that NOC1 overexpression did not significantly change the morphology or the size of cells, that developed at similar rate as control cells expressing GFP (Figure 6A-B, J). Instead, NOCs downregulation caused a significant reduction in the number of smaller clones, and cells exhibited a morphology that was reminiscent of dying cells (Figure 6C-E, J). A deeper analysis showed that *NOC1-RNAi* clones also induced at 48 hours AEL were not detected when analyzed at 90 hrs. AEL, while control GFP clones reached the size of about 120 cells/clone (Figure 6F-G, H). Only when recombination of *NOC1-RNAi* was induced at 72 AEL we were able to score few clones that were still significantly smaller than control and partially rescued when the inhibitor of caspase P35 was co-expressed. The size of these clones was overall 15% the size of wild type GFP clones, considered 100% (Figure 6K-L, I), and co-expression of P35 was able to partially rescue *NOC1-RNAi* clonal-size up to 60% (Figure 6M, I). These results suggest that cells with reduced NOC1 died and might be eliminated by neighboring cells by cell competition, a mechanism already described for ribosomal proteins of the *Minute* family and for nucleolar components regulating rRNA and ribosome maturation. To understand if *NOC1-RNAi* is involved in cell competition we induced *NOC1-RNAi* clones in animals heterozygotes for *Minute (3)66D* that carries a mutation in the gene encoding the ribosomal protein RpL14 (Saeboe-Larssen et al., 1997). These experiments show that *NOC1-RNAi* clones are partially rescued when induced in a *Minute* background both in their total number and size (Figure 6N-O), suggesting that NOC1 reduction activates or is part of the mechanisms regulating *Minute*-induced cell competition.

**Figure 6:**
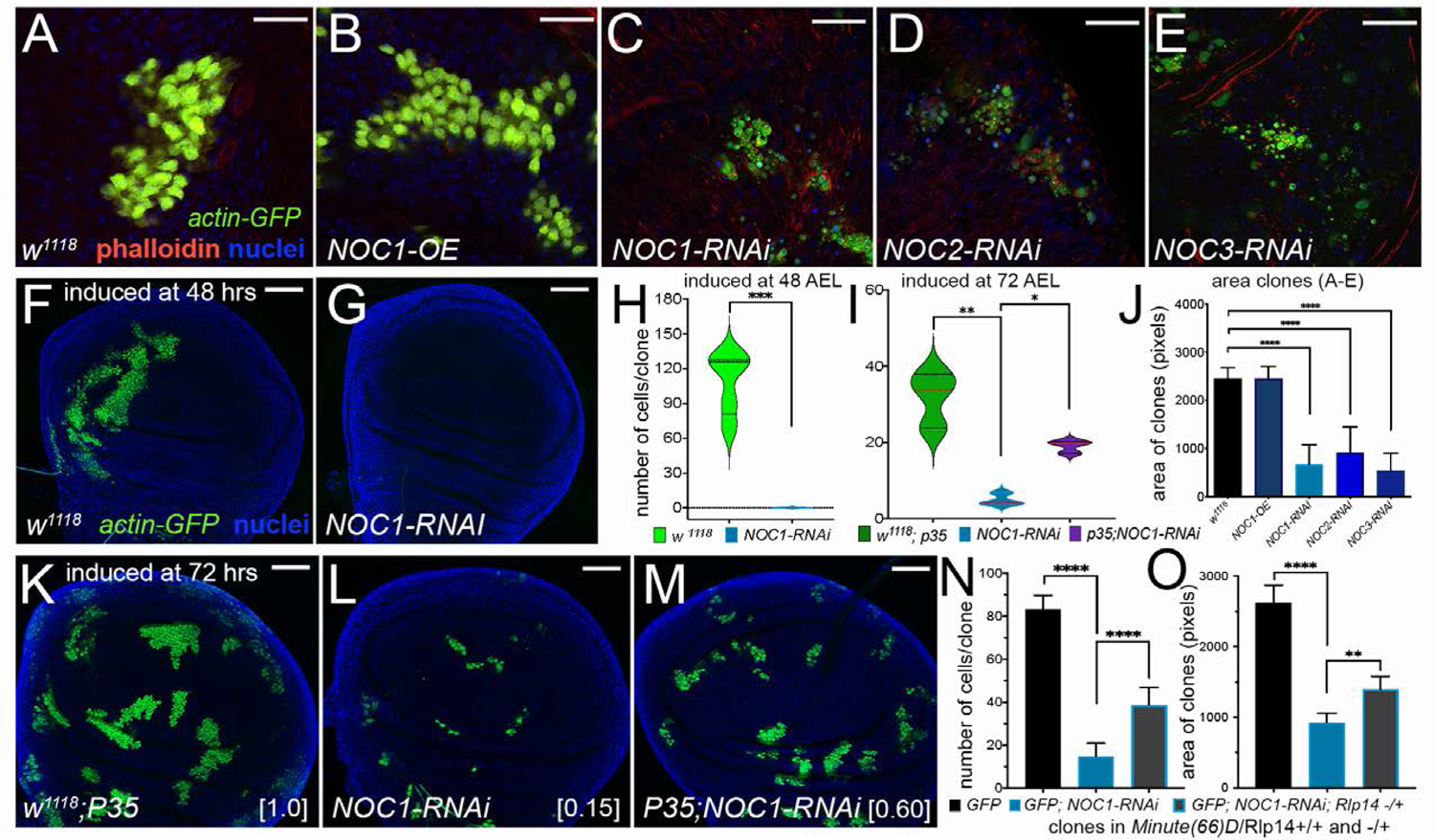
Reduction of NOC1, NOC2 and NOC3 in cells of the wing imaginal disc induces growth defects that are rescued by co-expressing P35 and in a *Minute (3)66D/+* heterozygous background. (A-E) Confocal images of *actin-*flip-out clones analyzed in the wing imaginal discs, expressing nuclear GFP and the indicated transgenes. Phalloidin-TR was used to mark the cell membranes (red) and Hoechst for the nuclei (blue). (J) Quantification of clonal size was performed by measuring the area marked by phalloidin; area is shown in pixels. At least 15 animals from each genotype were used; the scale bar represents 20 μm. (F, G) Confocal images of wing imaginal discs showing *actin-*flip-out clones expressing GFP alone (F) or co-expressing *NOC1-RNAi* (G). Clones were induced at 48 hours AEL; the scale bar represents 50 μm. (H-I) Quantification of the number of cells in each clone was analyzed at 120 hours AEL using GFP as marker. (K-M) Photographs of *actin*-flip-out clones in wing discs expressing GFP along with the inhibitor of caspase P35 (K), or with *NOC1-RNAi* (L) or co-expressing *NOC1-RNAi* together with P35 (M); clones were induced at 72 hours AEL. (I) The number of cells in each clone from K, L and M was quantified at 120 hours AEL. The total number of analyzed clones is indicated in parenthesis: *w^1118^* + P35 (72, [1.0]), *NOC1-RNAi* alone (66, [0.15]), and *NOC1-RNAi* + p35 (81, [0.60]). The numbers in square brackets represent the relative size of clones (average) compared to that from control, considered equal to 1. (N-O) Analysis cell number and clonal size of *NOC1-RNAi* clones induced in Rp+/- a Rp+/+ background using the *Minute(3)66D/+* line that carries a mutation in the Rpl14 protein. (N) Quantification of the number of clones in each disc. (O) Clonal size showing that defects of *NOC1-RNAi* cells are partially rescued when clones are grown in the *Minute(3)66D/+* (Rp-/+) background. Data in H were calculated using Student’s *t* test. The asterisks in the graphs in I, J, N and O represent the *p-*values from one-way analysis of variance (ANOVA) with Tukey multiple comparisons * = *p* < 0.05, ** = *p* < 0.01, *** = *p* < 0.001 and **** = *p* < 0.0001, and the error bars indicate the standard deviations. In figures F, G, K-M, Hoechst was used for staining the nuclei.

### Reduction of NOC1 in cells of the imaginal discs induces the pro-apoptotic gene *XRP1* and eiger/JNK signaling with the upregulation of DILP8 that delays development

We further characterized the mechanisms underlining NOC1-RNAi-induced apoptosis in the epithelial cells of the wing imaginal discs. Since previous analysis using *engrailed-Gal4* resulted in small and undeveloped imaginal discs, we switched to using the *MS1096* dorsal wing promoter (Capdevila and Guerrero, 1994) with which overexpressing NOC1 or its reduction did not significantly affected the morphology of the discs. The discs developed at similar rate as WT and showed the physiological Wingless pattern of expression at 96 hrs. AEL. (Figure 7A-C). However, when we deeper analyzed the timing of larval development, we observed that NOC1-RNAi larvae were delayed and reached pupariation in about 48 hrs. of difference with respect to control or NOC1-OE animals (Figure 7J, and Table I). In addition, NOC1-RNAi animals developed to a smaller size than control that reached a significant difference at 120hrs AEL (Figure 7G). Of note that reduction of NOC1 using the RNAi line on Chr. III resulted in pupal lethality, while using the line on Chr. II showed <10% of animals that hatched as small adults. These animals also presented morphological defects in the dorsal side of their wings, where *MS1096-Gal4* is expressed (Figure 7E and H). The combination of developmental delay and apoptosis prompted us to check if the reduction of NOC1 upregulates DILP8, that is normally secreted in response to cell death and tissue damage. Indeed, we found *DILP8* mRNA significantly increased in these animals (*p*<0.01), whereas its level was not changed upon NOC1 overexpression (Figure 7I).

**Figure 7:**
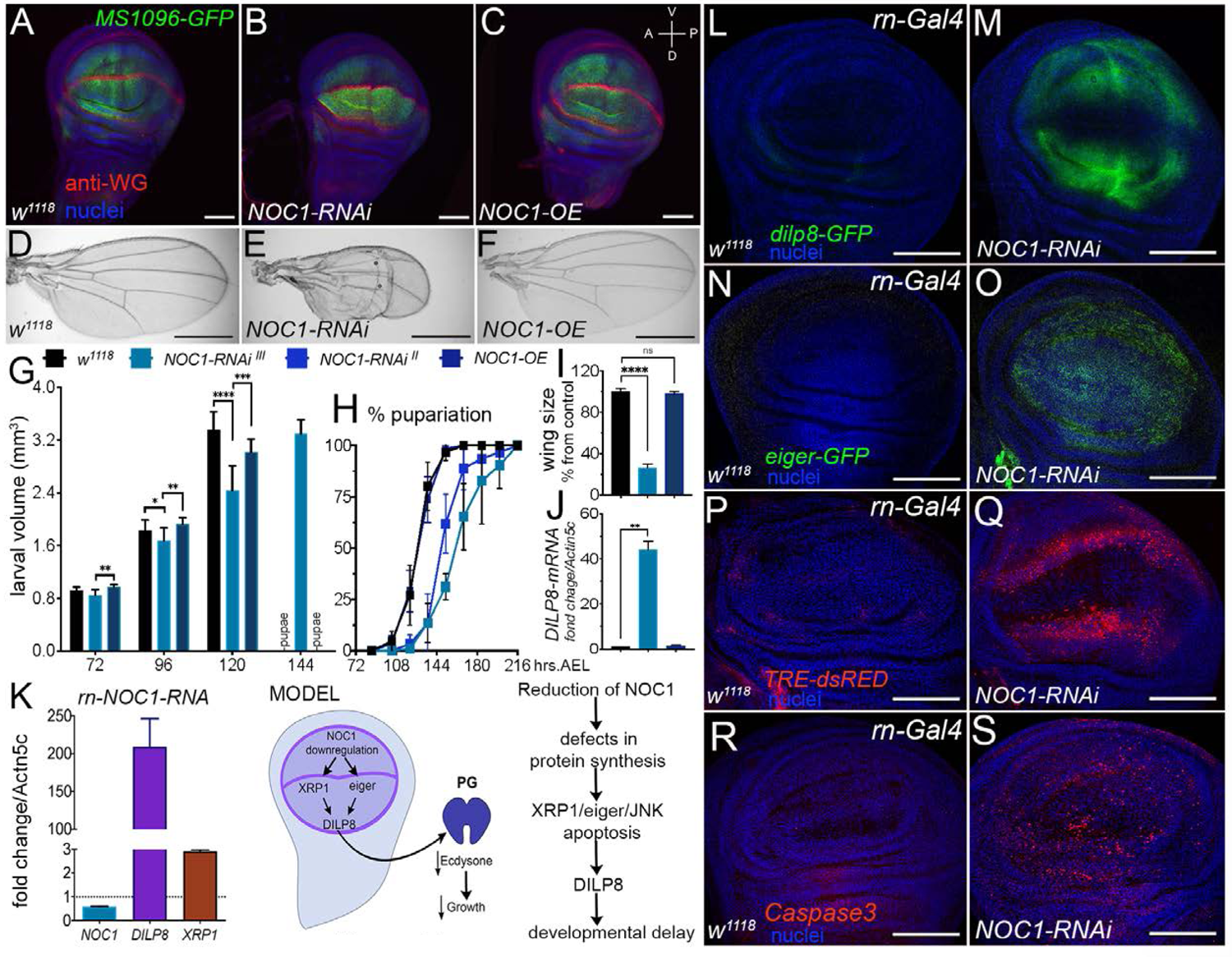
Reduction of NOC1 in cells of the wing imaginal disc induces Xrp1 and eiger resulting in apoptosis and DILP8-induced developmental delay. (A-C) Confocal images of wing imaginal discs expressing the indicated transgenes using the *MS1096-GFP* wing-driver. Wingless (WG) expression is visualized using anti-WG antibodies (in red), nuclei are stained with Hoechst (in blue); the scale bar represents 40 μm. (D-F) Photographs of wings from three days old males of the indicated genotype; the scale bar represents 1 mm. (G) Larval volume of animals expressing the indicated transgenes using the *MS1096*-driver was measured at the indicated time after egg laying (AEL) until pupariation. The graph is representative of one of three experiments using at least ten animals for each point and genotype, the asterisks represent the *p-*values from one-way (ANOVA) with Tukey multiple comparisons * = *p* < 0.05, ** = *p* < 0.01, *** = *p* < 0.001and **** = *p* < 0.0001, the error bars indicate the standard deviations. (H) Curves representing the % of pupariation of animals at the indicated genotypes. A significant delay in pupariation (one-way ANOVA, p < 0,0001) is visible in animals in which *NOC1* is reduced using *MS1096-Gal4* with both the RNAi lines. Data are expressed as % of pupariation over the total number of pupae of the same genotype, and the error bars indicate the standard deviations from five independent experiments. (I) Quantification of the area of the wings in male adult animals of the indicated genotypes at four days after eclosion. Data are expressed as % from control *w^1118^*. (J) qRT-PCR showing the level of *DILP8* mRNA in wing imaginal discs of third instar larvae overexpressing or downregulating *NOC1* using the *MS1096* promoter; *actin5C mRNA* was used as control. (K) qRT-PCR showing the level of *DILP8* mRNA and *Xrp1* mRNA in wing imaginal discs in which NOC1 level was reduced using NOC1-RNAi under the *rotund-Gal4* promoter. *Actin5C* mRNA was used as control. (L-S) Confocal images of wing imaginal discs from third instar larvae expressing *DILP8-GFP* (L-M), *eiger-GFP* (N-O) and TRE-dsRED (P-Q) reporters and stained for anti-Caspase3 (R-S); NOC1 was reduced using RNAi under the control of the *rotund-Gal4* driver. The scale bars represent 100 μm. MODEL: NOC1 is necessary for proper rRNA processing. Its reduction decreases protein synthesis and induces a nucleolar stress resulting in apoptosis. This event is accompanied by the upregulation of the pro-apoptotic genes *eiger* and Xrp1, both described to converge on the regulation of DILP8 that is induced to reduces ecdysone to delays animal development.

To better analyze at cellular level the mechanism underlying DILP8 upregulation, we reduced NOC1 using the wing specific *rotund-Gal4* promoter. These experiments confirmed that in cells where NOC1 was reduced the level of *dilp8-GFP^M100727^* was increased (Figure 7M), reflecting the robust increment of *DILP8* mRNA detected in the wing imaginal discs (Figure 7K) (similar results were obtained using the *MS1096* promoter - not shown). Concomitantly we found an upregulation of the pro-apoptotic gene *eiger,* visualized using its reporter *eiger-GFP^fTRG^* (Figure 7O), that was accompanied with an activation of JNK signaling indicated by the increase in *TRE-dsRED* (Figure 7Q). In agreement with these results, anti-Caspase3 staining indicates that apoptosis was significantly increased in cells with reduced NOC1 (Figure 7S). Cells undergoing proteotoxic stress are subjected to elimination by cell competition with a mechanism that depends on Xrp1, a transcription factor upregulated in Rp-/+ conditions. Notably, we found that these cells transcriptionally upregulate Xrp1 upon NOC1 reduction (Figure 7K). These data strongly suggest that cells with reduced NOC1 undergo proteotoxic stress with upregulation of *eiger* and *Xrp1,* causing cell damage that activates DILP8/Lgr3 compensatory mechanism responsible for the developmental delay observed in *NOC1-RNAi* animals.

### NOC1 CRISPR mutation affects animal growth and phenocopies NOC1 downregulation induced by RNA interference in the wing disc

To develop genomic *NOC1* mutants, we induced site specific mutations with the CRISPR-Cas9 system, using the line *sgRNA^CG7839^* from Boutros’s laboratory (Port et al., 2020). To understand if the reduction of NOC1 in this system phenocopied our data with *engrailed*-Gal4, we used a line that carries the *hedgehog-Gal4* to drive *UAS-Cas9* to express *sgRNA^CG7839^* in the posterior compartment of the wing disc. As shown in Figure 8, driving mutations of NOC1 using *hedgehog-Gal4* compromised and reduced the development of the posterior compartment of the wing disc with a similar extent to that observed with *engrailed-NOC1-RNAi* (Figure 8B and F). To compare the efficiency of the two systems, we analyzed the total area of imaginal discs and the ratio between the area of the posterior compartment (marked by co-expression of GFP) and the anterior from animals at 120 hours AEL. This analysis showed that reduction of NOC1 using *sgRNA^CG7839^* affected the total area of the discs and the ratio between the posterior and the anterior compartments (Figure 8C and D). These data resembled that obtained using NOC1-RNAi expressed under the *engrailed* promoter (Figure 8G and H). To introduce NOC1 mutations in the germ line, we used the *nos-Gal4, UAS-Cas9* line crossed with *sgRNA^CG7839^*. Sequencing analysis of 30 NOC1 heterozygous lines revealed the presence of missense mutations in the NOC1 gene in two lines, which encoded for very short polypeptides of 30 and 29 amino acids in length in NOC1-mut12 and NOC1-mut14, respectively (Figure 8K). Moreover, the phenotypic analysis of these two homozygous NOC1 mutants showed a robust growth defect at the larval stage (Figure 8I and J), recapitulating the phenotype described in the *actin-NOC1-RNAi* larvae (Figure 1B).

**Figure 8:**
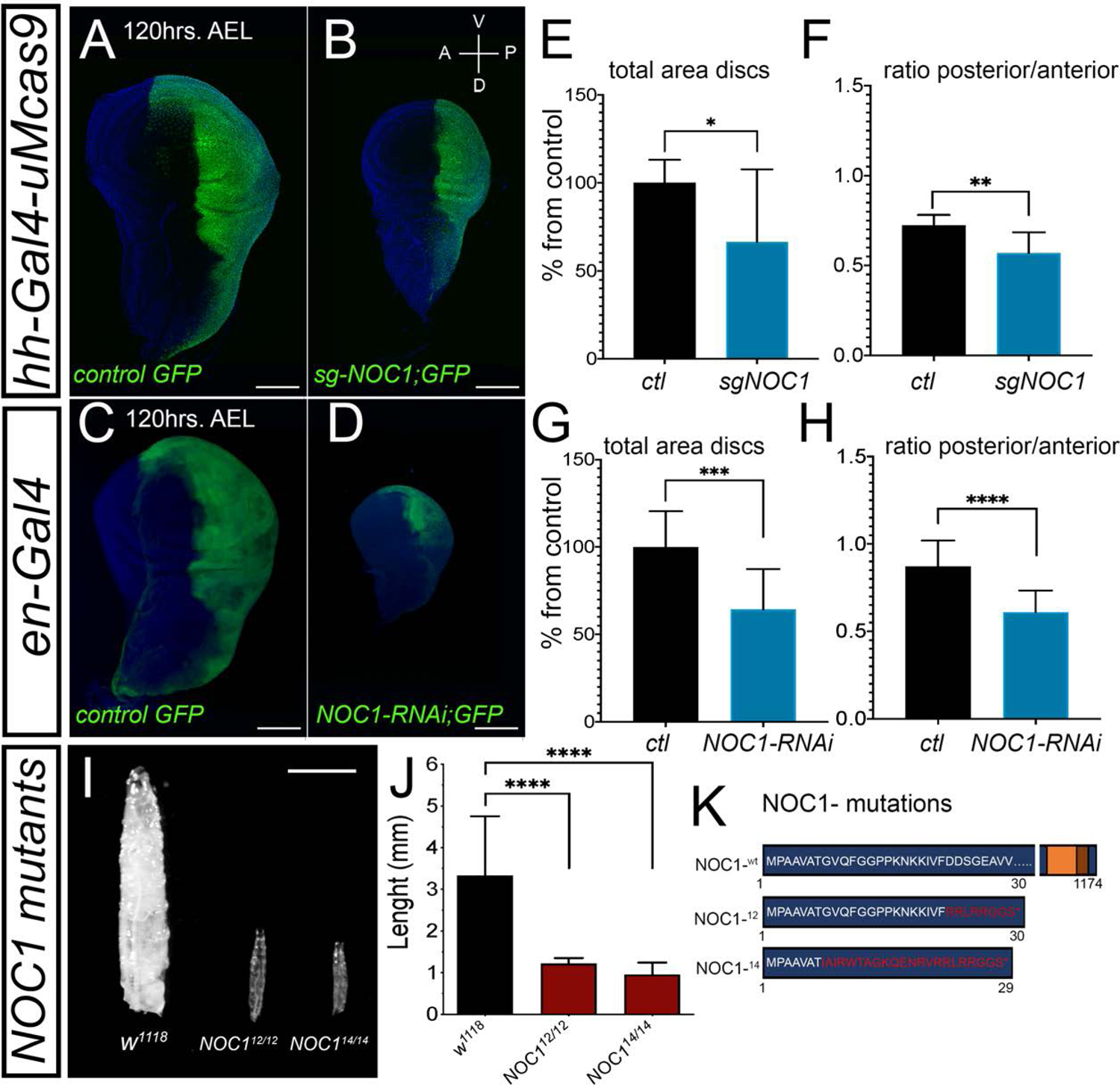
Targeted NOC1 CRISPR-mutation phenocopies the effect of NOC1 downregulation using RNA interference. (A-B) Confocal images of wing imaginal discs from larvae at 120 hours AEL, expressing *hedgehog-Gal4-GFP (hh-Gal4)* along with *uMcas9* (A) or with *uMcas9* and the *sg-RNA^CG7839^*(B), to conditionally knockout *NOC1* in cells of the posterior compartment of the wing disc marked by GFP expression. (E-F) Confocal images of wing imaginal discs in which *engrailed (en)-Gal4* was used to drive the expression in the posterior compartment of GFP alone (E) or together with *NOC1-RNAi ^II^* (F); the scare bars represents 50 μm. (C-D) Quantification of the total area of the wing imaginal discs (C) and of the posterior/anterior compartments ratio (D) from A and B. (G-H) Quantification of the total area of the wing imaginal discs (G) and of the posterior/anterior compartments ratio (H) from E and F. Data in C-D and G-H are representative of one of three experiments using ten or more animal for each genotype; the asterisks represent the *p-*values from Student’s *t*-test * = *p* < 0.05, ** = *p* < 0.01, *** = *p* < 0.001and **** = *p* < 0.0001, and the error bars indicate the standard deviations. (I) Photographs of third instar control *w^1118^* larvae and of *NOC1^12/12^* and *NOC1^14/14^* homozygous mutants taken at 120 hours AEL; the scale bar represents 1 mm. (J) Larval length of *NOC1^12/12^* and *NOC1^14/14^* homozygous mutants (*p* < 0.0001, one-way ANOVA). More than 10 animals for each genotype were analyzed. (K) Schematic diagram of NOC1^12/12^ and NOC1^14/14^ mutants that should encode for short polypeptides of 30 and 29 amino acids, respectively.

## Discussion

Here, we show that *Drosophila’s* homologues NOC1, NOC2 and NOC3 are necessary for proper animal development as their ubiquitous reduction results in growth defect and larval lethality (Table 1 and Figure 1A). NOC1 ubiquitous overexpression does not apparently affect larval transition, however these animals die at pupal transition, a condition that is rescued by co-expression of NOC1-RNAi (Figure 1D-H). These data suggest that NOC1 levels must be tightly controlled, as either its reduction or overexpression is detrimental for the cells. As demonstrated for its yeast’ homologue, NOC1 function in *Drosophila* is not redundant and its overexpression does not compensate for the loss of *NOC2* and *NOC3*. The reason for this unicity may be that NOC proteins work together in functional heterodimers (NOC1/NOC2 and NOC2/NOC3) necessary to control rRNA processing and nascent 60S ribosomal subunits (Edskes et al., 1998; Hierlmeier et al., 2013; Milkereit et al., 2001). Indeed, it was demonstrated in yeast that NOC1/NOC2 complex regulates the activity of the assembling factor Rpr5 to control rRNA cleavage at the internal transcribed spacers ITS1 and ITS2 sequences, to ensure the maturation of stoichiometric amount of 40S and 60S subunits (Khoshnevis et al., 2019). This function is likely to be conserved also in flies. In fact, our results show that reduction of NOC1 induces the accumulation of the intermediate ITS1 and ITS2 immature forms of rRNAs. Moreover, we observed a reduction in the relative abundance of 18S and 28S rRNAs (Figure 2F), suggesting that NOC1 is required for proper rRNA processing and ribosome maturation (Milkereit et al., 2001). In line with this hypothesis, we demonstrated that NOC1 reduction results in a strong decrease in ribosome abundance and assembling, also accompanied by a strong reduction of the 80S and the polysome profiling (Figure 2C). In addition, we also observed a mild accumulation of the 40S and 60S subunits (Figure 2D and E), perhaps suggesting that the mature 80S ribosome might be unstable in NOC1-RNAi animals and that a small percentage can disassemble in the two subunits, leading to the observed small increase. In addition, since NOC1 was identified as a predicted transcription factor (Kudron et al., 2018; Neumuller et al., 2013; Port et al., 2020), and because reduction of NOC1 results to a robust decrease in global protein synthesis (Figure 2G-H), we cannot exclude that specific factors involved in the 80S assembling can be reduced or missing in these animals.

Analysis of protein-protein interaction using STRING indicates that CG7838/NOC1 may act in a complex with other nucleolar proteins (Supplementary Figure S2). Indeed, here we showed that NOC1 localizes in the nucleolus with fibrillarin (Figure 1I, K-N). However, NOC1 overexpression also results in the formation of large rounded nuclear structures, reduced when its expression is silenced with NOC1-RNAi (Figure 1L and N). Interestingly, similar structures have been shown for CEBPz, the human homologue of NOC1 visible in images from “The Human Protein Atlas”. CEBPz (also called CBF2, CTF2) (OMIM-612828) is a transcription factor member of the CAAT-Binding proteins, involved in the complex of H*sp70* activation (Lum et al., 1990) and found upregulated in tumors, particularly in cells from patients with Acute Myeloid Leukemia (AML) (Herold et al., 2014). The presence also in NOC1 of the conserved CBP domain (Figure 1A) suggests that it may act as transcription factor, hypothesis corroborated by metadata in *Drosophila* (CHIP-Seq and genetic screens) that demonstrate how its expression is associated to promoter regions of genes important for the regulation of nucleolar activity and of ribosomal proteins (Neumuller et al., 2013; Shokri et al., 2019). This observation opens the possibility that NOC1 may control ribosome biogenesis through alternative mechanisms in addition to its control on rRNA transport and maturation. Moreover, we believe this function may be conserved for CEBPz, since in our bioinformatic analysis we identified nucleolar components and ribosomal proteins being upregulated in liver and breast tumors with overexpression of CEBPz (Supplementary Figure S6). Interestingly, misexpression of some of these targets, like Rpl5 and Rpl35a, have been associated to ribosomopathies suggesting the possibility that CEBPz could also contribute to tumorigenesis in these genetic diseases (Mills and Green, 2017; Narla and Ebert, 2010).

To better characterize NOC1 functions *in vivo* we modulated its expression in organs that are relevant for *Drosophila* physiology, such as the prothoracic gland (PG), the fat body (FB) and the wing imaginal discs.

### Prothoracic gland (PG)

While the overexpression of NOC1 in the prothoracic gland (PG) does not affect development, its reduction significantly decreases ecdysone production, as shown by *E74b* mRNA levels (Figure 4D). This reduction is significant at 12 days AEL and occurs concomitantly with the reduction of the PG size (Figure 4A-C). Consequently, these animals are developmentally delayed and do not undergo pupariation but rather continue to wander until they die at about 20 days AEL. These animals feed constantly and increase their size, accumulating fats and sugars in the fat body’s cells that augment their size (Figure 4E-F). We previously described the presence of hemocytes (macrophage-like cells) infiltrating the fat body of these animals, a condition accompanied with an increase in JNK signaling and ROS (Radical Oxygen Species), likely released by the fat cells under stress condition (Valenza et al., 2018). Interestingly, this inter-cellular event recapitulates the chronic low-grade inflammation, or Adipocyte Tissue Macrophage (ATM), a pathology associated to adipose tissue in obese people (Horng and Hotamisligil, 2011) that represents the consequence of impaired lipid metabolism.

### Fat body

Reduction of NOC1, 2 or 3 in the fat body results in smaller and fewer cells while reduction of NOC1 in the whole organ inhibits its function (Figure 5C-E and N). The fat body regulates animal growth by sensing amino acids concentrations in the hemolymph and remotely controls the release of DILP2, 3 and 5 from the Insulin Producing Cells (IPCs) (Andersen et al., 2013; Geminard et al., 2009; Hyun, 2018). The fat body also functions as storage of nutrients (fats and sugars) necessary during the catabolic process of autophagy that allows animals to survive metamorphosis (Rusten et al., 2004; Scott et al., 2004). When nutrients are limited, larvae delay their developmental to accumulate fats and sugars until reaching their critical size, that ensure them to progress through metamorphosis (Hironaka et al., 2019; Texada et al., 2020). NOC1 downregulation in the fat alters their ability to store nutrients, and larvae proceed poorly through development (Figure 5G). In addition, these animals show DILP2 accumulation in the IPCs even in normal feeding conditions (Figure 5Q), indicating that the remote signals responsible for DILPs release are greatly reduced, phenocopying animals in starvation or with reduced levels of MYC in fat cells (Geminard et al., 2009; Parisi et al., 2013). Interestingly, we also observed that *Cg-NOC1-RNAi* animals accumulate an abnormal amount of fats in non-metabolic organs such as gut, brain, and imaginal discs (Figure 5O-P). This finding suggests that these animals are subjected to inter-organ dyslipidemia, a mechanism of lipids transport activated when the fat body is impaired, which triggers non-autonomous signals to induce other organs to store fats. Interestingly, this condition recapitulates dyslipidemia in humans, where the compromised adipose tissue releases lipoproteins of the APO family inducing fat accumulation in organs (Pirillo et al., 2021). Notably, a similar condition was described also in flies for mutations in members of the *APOE* family (Palm et al., 2012), outlining how the mechanisms controlling the inter-organ fat metabolism are conserved among species.

### Wing imaginal discs

NOC1 depletion in clones analyzed in the wing imaginal discs triggers their elimination by apoptosis (Figure 6K-M and Figure 7R-S). This event is partially rescued when clones are induced in the hypomorphic background of the *Minute(3)66D/+* mutation (Saeboe-Larssen et al., 1997) (Figure 6N-O). These cells also upregulate the pro-apoptotic gene Xrp1, recently showed to be responsible for controlling translation and indirectly for regulation of cell competition upon proteotoxic stress (Baillon et al., 2018; Baumgartner et al., 2021; Kiparaki et al., 2022). Reduction of NOC1 in the wing imaginal disc prolongs larval development (Figure 7G-J) with upregulation of DILP8 (Figure 7I and M) normally induced by cellular damage and apoptosis. The fact that NOC1-RNAi cells upregulate, in addition to Xrp1, *eiger* (Figure 7N-O), another pro-apoptotic gene and member of the TNFα family and activate JNK pathway (Figure 7P-Q), suggests that different mechanisms are converging in these cells to induce apoptosis and DILP8 upregulation. We are currently performing genetic epistasis to define in deep the relation between eiger signaling in NOC1-RNAi cells and how this is linked to Xrp1 transcriptional upregulation in response to nucleolar stress and DILP8 upregulation. Our hypothesis is that eiger may act in parallel or be regulated by other components that link defects in protein synthesis to apoptosis in cell competition, including RpS3 (Baumgartner et al., 2021) and RpS12 (Akai et al., 2021; Ji et al., 2019) ribosomal proteins, p53 (Sanchez et al., 2019) and its target Xrp1 (Boulan et al., 2019).

Collectively, here we described a new role for the nucleolar proteins NOC1, NOC2 and NOC3 in the control of animal growth in *Drosophila*. We particularly showed the relevance of NOC1 in promoting nucleolar stress and apoptosis, both leading cause of tumor formation (Penzo et al., 2019; Quin et al., 2014). Our data also support the potential role of NOC1 human homologue CEBPz in the context of tumorigenesis, which has not been elucidated yet. Indeed, mutations in CEBPz are described in >1.5% of tumors of epithelial origins (cBioPortal), suggesting that it may have a role in contributing to the signals that trigger proteotoxic stress associated with tumorigenesis (Mills and Green, 2017; Narla and Ebert, 2010). CEBPz activity was also associated to regulation of cellular growth together with the METTL3-METTL14 methyltransferase complex (Barbieri et al., 2017) and to the regulation of H3K9m3 histone methylation in response to srHC (sonication-resistant heterochromatin), highlighting other roles for this transcription factor in RNA and heterochromatin modifications (McCarthy et al., 2021).

## Materials and Methods

### Drosophila husbandry and lines

Animals were raised at low density in vials containing standard fly food, composed of 9g/L agar, 75 g/L corn flour, 50 g/L fresh yeast, 30g/L yeast extract, 50 g/L white sugar and 30 mL/L molasses, along with nipagine (in ethanol) and propionic acid. The crosses and flies used for the experiments are kept at 25°C, unless otherwise stated. The following fly lines were used: *GMR-Gal4* (Parisi et al., 2011), *tub>y+>Gal4; ey-flp* (Bellosta et al., 2005), *P0206-GFP-Gal4* (Valenza et al., 2018), the fat body specific promoter *FB-Gal4* (kind gift from Ines Anderl, University of Tampere, Finland,) *rotund-Gal4* (kind gift from Hugo Stocker, ETH Zurich, CH), *actin-Gal4,GFP/Gla*,*Bla* (kind gift from Daniela Grifoni University of l’ Aquila, IT), *yw; Actin>CD2>Gal4,GFP/TM6b* (kind gift from Bruce Edgar, University of Boulder, CO), *MS1096-Gal4* (kind gift from Erika Bach, NYU, USA), *Minute(3)66D/+ (Saeboe-Larssen et al., 1997), engrailed-Gal4,GFP* and *actin-Gal4, GFP; tub-Gal80ts* from the lab. The following stocks were obtained from the Bloomington *Drosophila* Stock Center: *Cg-Gal4.A2* (B7011), *elav-Gal4* (B458), *UAS-CG7839-RNAi (B25992*), *UAS-CG9246-RNAi* (B50907), *UAS-CG1234-RNAi* (B61872), *CG7839-GFP.FTPD* (B51967), *DILP8-GFPMI00727* (B33079); and from the Vienna *Drosophila* Resource Center: *UAS-CG7839-RNAi* (v12691), *sgRNA^CG7839^-CFDlib01132* (v341898), *hh-Gal4;uMCas9* (v340019), *w^1118^* (v60000), *eiger-GFP*-*2XTY1-SGFP-V5-preTEV-BLRP-3XFLAG* (v318615); and from FlyORF (ZH) the line *UAS-CG7839*-*3xHA* (F001775).

### Measurement of larval length and volume

Larvae at the indicated stage of development and genotypes were anesthetized using freezing cold temperature, and pictures were taken using a Leica MZ16F stereomicroscope. Width and length were measured using a grid and volume was calculated by applying the formula in (Parisi et al., 2013).

### Quantitative RT-PCR

RNA extraction was performed using the RNeasy Mini Kit (Qiagen), following the manufacturer instructions. The isolated RNA was quantified with the Nanodrop2000. 1000 ng of total RNA were retrotranscribed into complementary DNA (cDNA) using the SuperScript IV VILO Master Mix (Invitrogen). The obtained cDNA was used for qRT-PCR using the SYBR Green PCR Kit (Qiagen). The assays were performed on a BioRad CFX96 machine and the analysis were done using Bio-Rad CFX Manager software. Transcript abundance was normalized using *actin5c*. The list of the primers is available in Supplementary Figure 6.

### Dissection and immunofluorescence

Larvae were collected at the third instar stage, dissected in 1x phosphate-buffered saline (PBS), and fixed for 30 minutes in 4% paraformaldehyde (PFA) at room temperature (RT). After 15 minutes of tissue permeabilization with 0.3% Triton X-100, samples were washed in PBS-0.04% Tween20 (PBST) and blocked in 1% bovine serum albumin (BSA) for 1 hour at RT. Samples were incubated overnight at 4°C with primary antibodies in 1% BSA and, after washing, with Alexa-Fluor-conjugated secondary antibodies 1:2000 in BSA. During washing in PBST nuclei were stained with Hoechst. Imaginal discs were dissected from the carcasses and mounted on slides with Vectashield. Imagines were acquired using a Leica SP5 and SP8 confocal microscope and assembled using Photoshop2020 from Creative Clouds. Primary antibodies used: rat anti-HA 1:1000 (Roche 3f10) and anti-fibrillarin (ABCAM ab4566), anti WG 1:100 (DSHB 4D4).

### Western blot

Proteins were extracted from third instar larvae collected in 250 ul of lysis buffer (50 mM Hepes pH 7.4, 250 mM NaCl, 1 mM EDTA, 1.5% Triton X-100) containing a cocktail of phosphatases and proteases inhibitors (Roche). Samples were run on a SDS-polyacrylamide gel and then transferred to a nitrocellulose membrane. After blocking with 5% non-fat milk in TBS-Tween, membranes were incubated with primary antibodies against puromycin 1:000 (clone 12D10 MABE343, Merk), mouse anti-HA 1:200 (supernatant Sigma HA7) and anti-actin 1:200 (DSHB 224-236-1), followed by incubation with HRP conjugated secondary antibodies (Santa Cruz Biotechnology), and signal was detected using ECL LiteAblot Plus (Euroclone) and the UVITec Alliance LD2.

### SUnSET assay

*UAS-NOC1-RNAi* was expressed ubiquitously in whole larvae using the *actin-Gal4* coupled with *tubulin-Gal80* temp sensitive allele to avoid early lethality. Crosses were kept at 18°C and when larvae reached second instar were switched to 30°C for 72 hours prior to dissection. At least seven third-instar larvae for each genotype were dissected in Schneider’s medium and then transferred to Eppendorf tubes containing complete medium with 10 % serum plus puromycin at 20 µg/ml (Invitrogen, Thermo Fisher Scientific). The samples were incubated for 40 or 60 minutes at room temperature, then recovered in 10% serum/media without puromycin for 30 minutes at room temperature. After the inverted larvae were snap frozen in liquid nitrogen for subsequent western blot analysis using anti-puromycin primary antibody.

### Polysome profiling

Cytoplasmic lysates were obtained from snap-frozen whole larvae pulverized using liquid nitrogen. After addition of lysis buffer and centrifugations for removal of the debris, cleared supernatants were loaded on a linear 10%–40% sucrose gradient and ultracentrifuged in a SW41Ti rotor (Beckman) for 1 hour and 30 min at 270,000 *g* at 4°C in a Beckman Optima LE-80K Ultracentrifuge. After ultracentrifugation, gradients were fractionated in 1 mL volume fractions with continuous monitoring of absorbance at 254 nm using an ISCO UA-6 UV detector. % of ribosomal subunits was calculated over the (40, 60, 80 and polysome) area of the same genotype.

### Generation of inducible flip-out clones and clonal analysis

Females, *yw; Actin>CD2>Gal4-GFP/TM6b* were crossed with males carrying the heat-shock *Flippase y^122^w* together with the relative UAS-transgenes. Animals were left laying eggs for 3-4 hours. Heat shock was performed on larvae at 48 or 72 hours after egg laying (AEL) for 15 min at 37°C. Larvae were dissected at 96 or at 120 hours AEL and mounted using MOWIOL. Images of clones expressing nuclear GFP were acquired using a LEICA SP8 confocal microscope. Quantification of the number of GFP-positive cells/clone in the wing imaginal discs was calculated from 5 confocal images for every genotype at 40x magnification maintaining constant acquisition parameters. Co-staining with phalloidin-Rhodamine (Invitrogen) was necessary in Figure 5 A-E to outline the cell membranes and DAPI for the nuclei.

### Imaging the adult compound eye and wings

Photographs of eyes of adult female expressing the indicated UAS-transgenes in the retina using the *GMR-Gal4 or tub>y+>Gal4* promoters were taken at 8 days after eclosion using a Leica stereomicroscope MZ16F at 4x magnification. 2-4-days old animals were fixed in a solution of 1:1 glycerol and ethanol. One wing was dissected from at least 10 animals and mounted on a slide in the same fixing solution. Images of each wing were taken using a Zeiss Axio Imager M2 microscope with a 1x magnification. Quantification of the area of each wing was performed on photographs using PhotoshopCS4.

### Fat body staining and cell size calculation

Fat bodies were dissected from larvae at 5 or 12 days AEL fixed in 4% PFA and counterstained with Nile Red (Sigma), Phalloidin-488 (Invitrogen) and Hoechst 33258 (Sigma). After washing with PBS, fat bodies were mounted onto slides with DABCO-Mowiol (Sigma-Aldrich) and images were acquired using LeicaSP5-LEICA, the area of adipose cells for each fat body was calculated with ImageJ software. Measurement of TAGs and visualization of lipids in the whole larvae was done as in (Parisi et al., 2013) using Nile Red staining. Dissected organs were mounted in DABCO-Mowiol and photographs were taken using a Zeiss Axio M2 Imager light microscope.

### Generation of CRISPR-Cas9 mutations of *Noc1/CG7839* and analysis of their function in the posterior compartment of wing imaginal disc

To target mutations in CG7839 into the germ line we crossed the line nos-Gal4VP16 UAS-uMCas9(attP40) with the line *gRNA* for *CG7839* (*vCFDlib01132*) from Boutros collection (Port et al., 2020). Out of 30 putative lines carrying potential NOC1 heterozygous mutations, we sequenced five lines and two of them contained indels that create nonsense mutations that translated *NOC1-mRNAs* into short NOC1 polypeptidic sequences of 30 amino acids (NOC1-mut^12^) and 29 amino acids (NOC1-mut^14^). Sequences of the primers used for the screening are in Supplementary Figure S7.

Phenotypic analysis of NOC1-mutant homozygous larvae was carried by leaving heterozygous *w^1118^; NOC1-mutant/TM6b* parents to lay eggs for 5 hrs. at 25°C in regular food. Homozygous not *tubby* larvae were scored, and pictures were taken at 8 days AEL. At this stage heterozygous *NOC1/TM6b* larvae were all pupae that hatched at the expected time.

Mutations of *CG7938* were targeted in the posterior compartment of the wing imaginal disc by using the line *UAS-uMCas9; hh-Gal4/TM6B* to spatially limit the transcription of Cas9 in the posterior region of the animal under *hh-Gal4* (Port et al., 2020). This line was crossed with that carrying the *sgRNA* for *CG7839* (*vCFDlib01132*) previously recombined with *UAS-GFP* to mark the posterior compartment. A line expressing only *UAS-GFP* was used as control. F1 animals were dissected at about 90 hours AEL and images of their wing imaginal discs were acquired using a confocal microscope (Leica SP8). Calculation of the size of the posterior compartment (GFP-positive) and the total area of the wing imaginal discs were performed using Adobe Photoshop (Creative Cloud). At least 8 animals from each genotype were used for the statistical analysis.

## Statistical Analysis

Student *t*-test analysis and analysis of the variance calculated using one-way ANOVA with Tukey multi-comparisons test were calculated using GraphPad-PRISM*8. p* values are indicated with asterisks ***\* =*** *p < 0.05, ** = p < 0.01, *** = p < 0.001, **** = p < 0.0001*, respectively.

## Acknowledgements

We thank Marcello Ceci for reading the manuscript and for the useful discussion, the Advanced Imaging Core Facility at CIBIO, the Vienna VDRC and Bloomington Stocks Centers and the DSHB for antibodies. Stocks obtained from the Bloomington Drosophila Stock Center (NIH P40OD018537) were used in this study. We apology in advance to any authors whose work has been omitted.

## Data availability

none

