## Supplementary material for "Reduction of nucleolar NOC1 accumulates pre-rRNAs and induces Xrp1 affecting growth and resulting in cell competition in *Drosophila*": suppl data

### Fig. S1.  NOC proteins alignments

#### NOC1

Query: CG7839-PA (IP14658p). Query ID: Q9VTE6 Length: 1174
Q03701 >CCAAT/enhancer-binding protein zeta [Homo sapiens] Sequence ID: NP_005751.2 Length: 1054
Range 1: 182 to 1019
Identities:272/922(30%), Positives:436/922(47%), Gaps:102/922(11%)

```
Query  239   KWYHV-----HPDYPSTDEVLDMKENDQLELYNLCKNSFEAEKITFNKRNPSDARWLQTA  293
             KWY +     +   P   +V+   +    +LY    N F+++    N +  + + W++
Sbjct  182   KWYDLEYSNEYSLKPQPQDVVSKYKTLAQKLYQHEINLFKSKT---NSQKGASSTWMKAI  238

Query  294   LHKGTAKDRANAGALLVTSNPLGNLEALSTLIGFCKISNKASNDVIAVLTDLWQEVL---  350
             +  GT  DR  A  LL+  + +  L+ + TL+   K   K S     +  D ++E+L
Sbjct  239   VSSGTLGDRMAAMILLIQDDAVHTLQFVETLVNLVK--KKGSKQQCLMALDTFKELLITD  296

Query  351   -LPPNRKLLAVHTRGADWKKLKKDENLRNEQKRRIYAYWHFESELKDQYHEFLKNVMQGL  409
              LP NRKL     R  D  KL++  +    + + R    W+FE +LK    EF++ +
Sbjct  297   LLPDNRKLRIFSQRPFD--KLEQLSSGNKDSRDRRLILWYFEHQLKHLVAEFVQVLETLS  354

Query  410   QTGQEHNKNSSIVSAARLLAYAPEKEQLLLTMLVNKLGDPIAKIASKALHHLSEVAQKHP  469
                    K  ++  A  LL   PE+E+ LL  +VNKLGDP  +IA+KA H L  +  KHP
Sbjct  355   HDTLVTTKTRALTVAHELLCNKPEEEKALLVQVVNKLGDPQNRIATKASHLLETLLCKHP  414

Query  470   NMCGVIVAEAEKLLFRNNISERAQHFALCFLSSIAPSG-RPEVCTKLVNICFALFKVLVQ  528
             NM GV+  E E+LLFR+NIS +AQ++A+CFL+ +A S    E+  KL+ + F  F+  V+
Sbjct  415   NMKGVVSGEVERLLFRSNISSKAQYYAICFLNQMALSHEESELANKLITVYFCFFRTCVK  474

Query  529   KGAVNNRTMQAILRCLQKAIVEAKPAKDSNGELLTKEMQDTIYRLVHLADIRVAVQTLGL  588
             K  V ++ + A+L  + +A    ++   D       +E  DT+++++H+ +   +VQ L L
Sbjct  475   KKDVESKMLSALLTGVNRAYPYSQTGDDK-----VREQIDTLFKVLHIVNFNTSVQALML  529

Query  589   LLQLVAVKTEKSDRFYNALYVKLLDLNLINVGSKTAAHLLHIVHRAIHIDNHVARAQAFV  648
             L Q++  +   SDR+Y ALY K+LD  L+    +  A  L++V++++  D  + R +AFV
Sbjct  530   LFQVMNSQQTISDRYYTALYRKMLDPGLMTCSKQ--AMFLNLVYKSLKADIVLRRVKAFV  587

Query  649   KRLLQLTLYAPPHIAAGCLIVIHKLLRMRRELIGGTGASEEVEEGSKVV-LPISADLDKF  707
             KRLLQ+T    P    G L ++ ++L+ +  L         E ++    +      D++KF
Sbjct  588   KRLLQVTCQQMPPFICGALYLVSEILKAKPGLRSQLDDHPESDDEENFIDANDDEDMEKF  647

Query  708   GSDDEEVYEDVKDEADDTKDSNPLEEKADNDVKSSASSWHHARVAATEAKVRDIDSCKYD  767
                D+E     K E ++T     +E K     K   +SW H         ++      KYD
Sbjct  648   TDADKETEIVKKLETEETVPETDVETK-----KPEVASWVHFDNLKGGKQLN-----KYD  697

Query  768   PYHRVPAFAGAAYALRHELLLLRQHYHPTVQVFAEQILQQSRIDYYGDPLRDFGLPHFLE  827
             P+ R P F GA      EL  L  H+HP+V +FA+ ILQ + I Y GDPL+DF L  FL+
Sbjct  698   PFSRNPLFCGAENTSLWELKKLSVHFHPSVALFAKTILQGNYIQYSGDPLQDFTLMRFLD  757

Query  828   RFAFKNPKKLEASQAAENATVAHKR-YMAHGARGRPVKS---LTK--ANCTEDEMFIFNF  881
             RF ++NPK  +  +  ++  +  KR +     R  PV S   L K  +    DE+F
Sbjct  758   RFVYRNPKPHKGKENTDSVVMQPKRKHFIKDIRHLPVNSKEFLAKEESQIPVDEVFF---  814

Query  882   LEHKRRQAEIVAQNKKQKEIKKDAAEEGDDGEAGEEYLKEGEVDDDEFEAYLDGYFGKKF  941
                  R  + VA  +KQ   K+DA EE   +    EE+ +  +  +D+          F
Sbjct  815   ----HRYYKKVAVKEKQ---KRDADEESIEDVDDEEFEELIDTFEDD----------NCF  857

Query  942   KEGVDEEQDEEELNFLQELGGEIKKDKSKDKKKKKQSDKAEDEMDDIDDDWGDDDLAEDE  1001
               G D+          +  G +          KK++  A+D   D D +  DD+L   +
Sbjct  858   SSGKDD----------MDFAGNV----------KKRTKGAKDNTLDEDSEGSDDELGNLD  897

Query  1002  DDEIEGEDQSDDETGSIDLQPLDDDDDDDDDDDDEGSISEGGPGDSDSSDAPESPDEEDD  1061
             DDE+      D+E   +D                    +GG       D  ES  E +
Sbjct  898   DDEVSLGSMDDEEFAEVD--------------------EDGGTFMDVLDDESESVPELEV  937

Query  1062  DDEDAPPRSKKSRKDSTDMVGGRSFAKTLKQSHDMSSLFAADDFSSLLEETAKVKGQGT  1121
               + +  +SK+   D  D  G     +  K++ + SSLF +A++F  LL+E    K
Sbjct  938   HSKVSTKKSKRKGTDDFDFAGSFQGPRKKKRNLNDSSLFVSAEEFGHLLDENMGSKFDNI  997

Query  1122  S-NAVFNKDKSSDKQLKWEENR  1142
               NA+ NKD +S KQL+WE  R
Sbjct  998   GMNAMANKDNASLKQLRWEAER  1019
```

#### NOC2:

Query: RecName: Full=Nucleolar complex protein 2 homolog; Short=Protein NOC2 homolog [Drosophila melanogaster] Query ID: Q9VIF0.1 Length: 766
nucleolar complex protein 2 homolog [Homo sapiens] Sequence ID: NP_056473.3 Length: 749 Sequence ID: Q9Y3T9.4
Range 1: 36 to 650
Identities:236/649(36%), Positives:385/649(59%), Gaps:45/649(6%)

```
Query  33   PQT-TSETKVTPRNPKQKVAEPVKNGKTTKKGFKKSHKEELEGLKDIDPEFYDFLKNNDK  91
            PQ  T E +   R+P +    P     + +KG    HK++L  LKD DPEFY FL+ ND+
Sbjct  36   PQAETREAREAARSPDKPGGSP---SASRRKGRASEHKDQLSRLKDRDPEFYKFLQENDQ  92

Query  92   KLLDFNLLDTDDDDDEEGDEEDKEDTVTKESKDDEDDEEKYHKPSKDLEVASDESDFEVD  151
             LL  N  D+D  ++EEG      D + + S++++  EE           +   ++   V
Sbjct  93   SLL--NFSDSDSSEEEEGPFHSLPDVLEEASEEEDGAEEGEDGDRVPRGLKGKKNSVPV-  149

Query  152  EEDDAAAGGIQKITLNLLHQWEQQLGQANISIDIVRKVIQAFNSALASISADGADGGENK  211
                         T+ ++ +W+Q   Q  ++  +  +V+QAF +A+A+   D      NK
Sbjct  150  -------------TVAMVERWKQAAKQ-RLTPKLFHEVVQAFRAAVATTRGDQESAEANK  195

Query  212  HNAAAFKVVGAAAFNGVIQLCVIHLQPAIIRLL----------GVRPNSSLPLHKHKKWV  261
                 F+V  +AAFN ++  C+  L   + +LL           ++P+SS PL     W
Sbjct  196  -----FQVTDSAAFNALVTFCIRDLIGCLQKLLFGKVAKDSSRMLQPSSS-PL-----WG  244

Query  262  KVRGCLRYYLTDLIRLVEQVSSPNILGVLLKHLHQMAGMVVPFSALGKTILKRLVVLWST  321
            K+R  ++ YL   I+LV  +S   +L  +L+H+  +     + F    + +LKR+V++WST
Sbjct  245  KLRVDIKAYLGSAIQLVSCLSETTVLAAVLRHISVLVPCFLTFPKQCRMLLKRMVIVWST  304

Query  322  GDETVRVLAFLCILKITRKQQATMLNHVLKAMYLAYVRNSKFVSPNTLPGINFMRRSLVE  381
            G+E++RVLAFL + ++ R ++ T L  VLK MY+ YVRN KF SP  LP I+FM+ +L E
Sbjct  305  GEESLRVLAFLVLSRVCRHKKDTFLGPVLKQMYITYVRNCKFTSPGALPFISFMQWTLTE  364

Query  382  MFALDLNVSYQHVFLYIRQLAIHLRNAVILKKKDSFQAVYNWQFINSLRLWADLLGASAN  441
            + AL+  V+YQH FLYIRQLAIHLRNA+  +KK++Q+VYNWQ+++ L LW  +L  +
Sbjct  365  LLALEPGVAYQHAFLYIRQLAIHLRNAMTTRKKETYQSVYNWQYVHCLFLWCRVLSTAGP  424

Query  442  KPQLQPLIYPLVTIATGVIRLIPTAQYFPLRFHCLQTLISLAKETNTYVPVLPLIVEVLK  501
               LQPL+YPL   +  G I+LIPTA+++PLR HC++ L  L+  +  ++PVLP I+E+ +
Sbjct  425  SEALQPLVYPLAQVIIGCIKLIPTARFYPLRMHCIRALTLLSGSSGAFIPVLPFILEMFQ  484

Query  502  SNTFNRKHSAVSMKPVQFTCVLRLNKGQLAENGFRDEVIEQVCGLLLEYLAHESTSLAFS  561
               FNRK   +S KP+ F+ +L+L+   L E  +RD ++EQ+  L LEYL  ++  + F
Sbjct  485  QVDFNRKPGRMSSKPINFSVILKLSNVNLQEKAYRDGLVEQLYDLTLEYLHSQAHCIGFP  544

Query  562  DLVVPTVMAIKTYLKECRNANYARKLKQLLEKIQESARFIEQQRGKSSVTFDIKDAQAVA  621
            +LV+P V+ +K++L+EC+ ANY R+++QLL K+QE++ +I  +R +  V+F + + QAV
Sbjct  545  ELVLPVVLQLKSFLRECKVANYCRQVQQLLGKVQENSAYICSRRQR--VSFGVSEQQAVE  602

Query  622  AWEQQLRLKRTPLDVYYASWLKTHETKKRRQAAHTDEINADYDVPKLKK  670
            AWE+  R + TPL +YY+ W K  + + + + +  + D + P++K+
Sbjct  603  AWEKLTREEGTPLTLYYSHWRKLRDREIQLEISGKERLE-DLNFPEIKR  650
```

#### NOC3:

Query: RecName: Full=Nucleolar complex protein 3 homolog; Full=Nucleolar complex-associated protein 3-like protein [Drosophila melanogaster] Query ID: Q9VI82.1 Length: 822
>nucleolar complex protein 3 homolog [Homo sapiens] Sequence ID: NP_071896.8 Length: 800 Sequence ID: Q8WTT2.1 Length: 800
Range 1: 204 to 778
Score:393 bits(1009), Expect:9e-129,
Identities:224/586(38%), Positives:350/586(59%), Gaps:19/586(3%)

```
Query  224  LIARQQEIERQKYRIGIICSGLLEKPEDKMRNFHALYELMDEINPASRQANLMAVRKLAI  283
            LI R+++++ +K  I  + S +L  PE+ ++    L  ++ E +P       + VRKL I
Sbjct  204  LIERKKKLQEKKMHIAALASAILSDPENNIKKLKELRSMLMEQDPDVA----VTVRKLVI  259

Query  284  ISVTEIFKDILPEYRVGQVDT--KMQTLRKATLDRVTFENALLQQFKKFLQKLEQITAQV  341
            +S+ E+FKDI P Y++  +    K    RK T     FE  L+ Q+K +L+ LEQ+
Sbjct  260  VSLMELFKDITPSYKIRPLTEAEKSTKTRKETQKLREFEEGLVSQYKFYLENLEQMVKDW  319

Query  342  NRRGGLRTPQTVKL------ATVAVQCMCDLLVAHPYFNYVQNIAQLLVYMLNCNYAEMR  395
             +R  L+    V L      A VAV+ +C+LLVA P+FN+  NI  L+V ++N     +
Sbjct  320  KQRK-LKKSNVVSLKAYKGLAEVAVKSLCELLVALPHFNFHNNIIVLIVPLMNDMSKLIS  378

Query  396  TAVHQCFRTVFSNDKRLEMTLFIVRRINHLIKTKQNNVHVECITCLMGLKIKNVNLDAEK  455
                +  + +F  DK  + +L +++ I+  +K +   V  E +    + L+IK V +  +
Sbjct  379  EMCCEAVKKLFKQDKLGQASLGVIKVISGFVKGRNYEVRPEMLKTFLCLRIKEVEVKKDT  438

Query  456  ENELKQKKLESHRQRLLSLSKKERKRRKKLTEVNRELEETRAEENKQAKHQKLTEIIKMV  515
            E+  K KK  + +++  SLS+ +RK +K   ++ REL E  A E+ + K +  TE + +V
Sbjct  439  EDINKPKKFMTFKEKRKSLSRMQRKWKKAEEKLERELREAEASESTEKKLKLHTETLNIV  498

Query  516  FTIYFRVLKNDPTSRVLSAILEGLAEFAHVINLDFFSDLIDVLNRILEDQDELGYRERLH  575
            F  YFR+LK    S +L A+LEGLA+FAH+IN++FF DL+ VL+ ++E  D L Y+E LH
Sbjct  499  FVTYFRILKKAQRSPLLPAVLEGLAKFAHLINVEFFDDLLVVLHTLIESGD-LSYQESLH  557

Query  576  CVQTIFVILSGQGEVLNIDPIRFYQHFYRNMLAVQAGKNHDDFAIILRTLDEVLVKRRRN  635
            CVQT F ILSGQG+VLNIDP++FY H Y+ +  + AG  ++   I+L+ LD +L KRR+
Sbjct  558  CVQTAFHILSGQGDVLNIDPLKFYTHLYKTLFKLHAGATNEGVEIVLQCLDVMLTKRRKQ  617

Query  636  MSQQRLMAFMKRLLTGSLHLLHNGTLATLGTIKQTFQLTSVLDNLLDTDTTIGSGRYDPE  695
            +SQQR +AF+KRL T +LH+L N ++  L T +         D LLD+++  GSG + PE
Sbjct  618  VSQQRALAFIKRLCTLALHVLPNSSIGILATTRILMHTFPKTDLLLDSESQ-GSGVFLPE  676

Query  696  LDDPEYCNAASTALYELALLARHYHPTVRRMAVHIAHGVPATGEGALPTEIGKLTSHELF  755
            LD+PEYCNA +TAL+EL  L RHYHP V+R A H+  G P+ G GAL  E+ + ++ ELF
Sbjct  677  LDEPEYCNAQNTALWELHALRRHYHPIVQRFAAHLIAGAPSEGSGALKPELSRRSATELF  736

Query  756  TQFDSTQMAFNPTIPLPKAGQPKLKRGKHLYIRSDFKQEYGKLLQQ  801
              +   +M FNP +   ++  PK+K GK L   S   ++  +L+++
Sbjct  737  EAYSMAEMTFNPPV---ESSNPKIK-GKFLQGDSFLNEDLNQLIKR  778
```

**Fig. S2. Protein-protein interaction network generated using STRING (Szklarczyk et al., 2019).** (A) Graphic representation and (B) predicted list of the functional partners of NOC1/CG7839, members of the interaction network.

# (A)

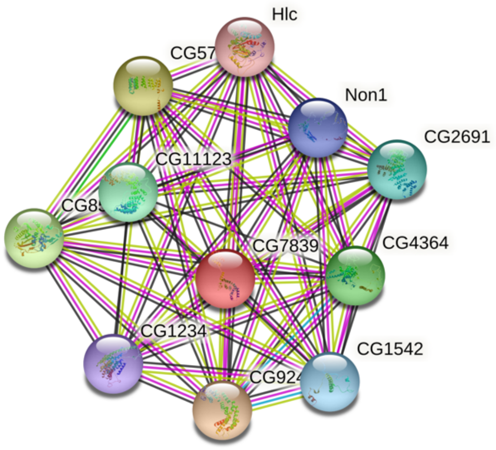

# (B)

**Your Input:**

| | |
|---|---|
| CG7839 | IP14658p; Sequence-specific DNA binding transcription factor activity. It is involved in the biological process described with: regulation of transcription, DNA-templated; neurogenesis (1174 aa) |

**Predicted Functional Partners:**

| Protein | Description | Neighborhood | Gene Fusion | Cooccurence | Coexpression | Experiments | Databases | Textmining | [Homology] | Score |
|---|---|---|---|---|---|---|---|---|---|---|
| CG9246 | Nucleolar complex protein 2 homolog; It is involved in the biological process described with: neurogenesis | | | | ● | ● | | ● | | 0.997 |
| CG5728 | LD41803p; mRNA binding. It is involved in the biological process described with: regulation of alternative mRNA splicing, via ... | | | | ● | ● | | ● | | 0.995 |
| CG8545 | LD11307p; RNA binding; S-adenosylmethionine-dependent methyltransferase activity. It is involved in the biological process d... | | | | ● | ● | | ● | | 0.995 |
| CG4364 | Pescadillo homolog; Required for maturation of ribosomal RNAs and formation of the large ribosomal subunit | | | | ● | ● | | ● | | 0.988 |
| CG11123 | RH42110p; RNA binding | | | | ● | ● | | ● | | 0.984 |
| CG2691 | RRP12-like protein; It is involved in the biological process described with: neuron projection morphogenesis | | | | ● | ● | | ● | | 0.982 |
| CG1542 | Probable rRNA-processing protein EBP2 homolog; Required for the processing of the 27S pre-rRNA | | | | ● | ● | | ● | | 0.982 |
| Non1 | Nucleolar GTP-binding protein 1; Involved in the biogenesis of the 60S ribosomal subunit (By similarity). Required for normal ... | | | | ● | ● | | ● | | 0.982 |
| CG1234 | annotation not available | | | | ● | ● | | ● | | 0.981 |
| Hlc | Helicase, isoform A; ATP binding; ATP-dependent RNA helicase activity; nucleic acid binding. It is involved in the biological pr... | | | | ● | ● | | ● | | 0.979 |

**Fig. S3. Length of larvae overexpressing *NOC1* or downregulating *NOC1, 2* and *3* in the whole animal using the *actin-Gal4* promoter.** Length was measured at 120 hrs AEL. The asterisks represent the *p*-values from one-way analysis of variance (ANOVA) with Tukey multiple comparisons ** = $p < 0.01$ and **** = $p < 0.0001$, and the error bars indicate the standard deviations for each genotype. In parenthesis is indicated the number of analyzed animals.

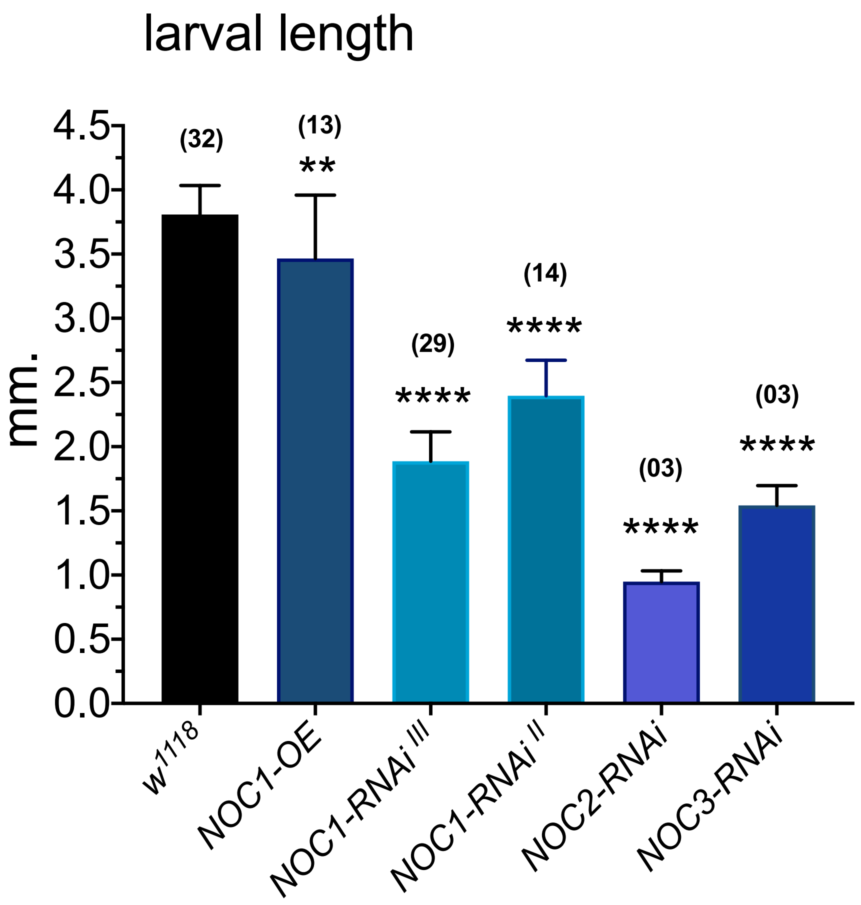

**Fig. S4: NOC1 localizes in the nucleolus in cells of the salivary glands.**

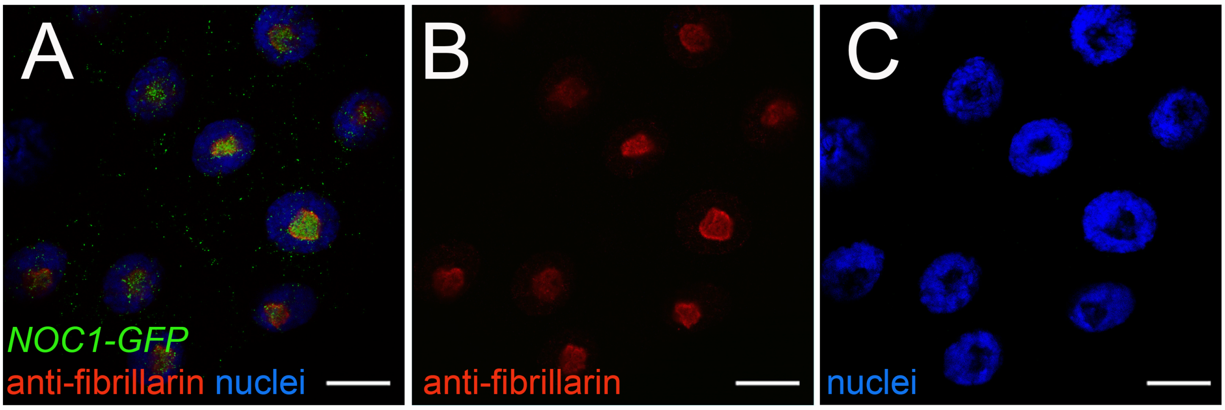

Analysis in cells in the salivary glands shows NOC1-GFP colocalization with fibrillarin in nucleoli, as demonstrated by co-immunostaining using anti-fibrillarin and anti-GFP antibodies. Nuclei were stained with Hoechst. The scale bars represent 20 μm.

**Fig. S5. qRT-PCR showing the relative amount of *NOC1 mRNA* upon RNA interference using two different RNAi lines.** *NOC1-RNAi* were ubiquitously expressed using the actin-Gal4 promoter. RNA was extracted from whole larvae.

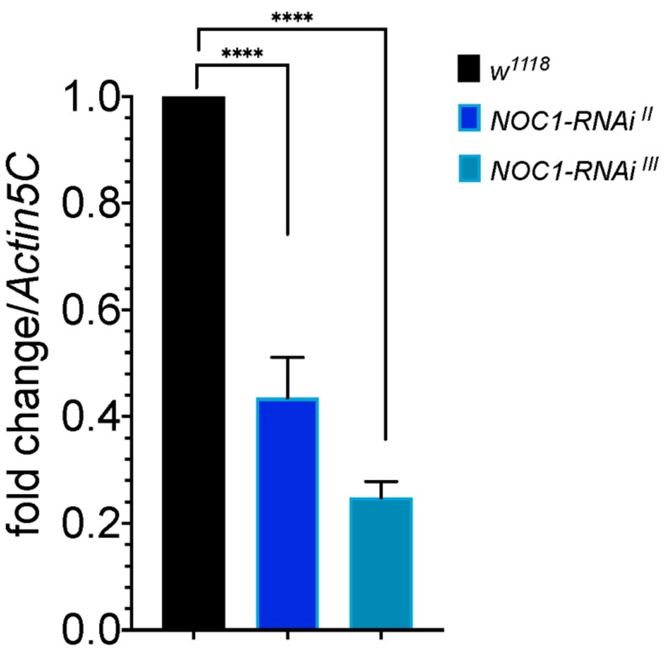

**Fig. S6: Selected list of potential targets of CEBPz involved in ribosomal biogenesis and nucleolar control.** Data are from TCGA datasets from cBio Cancer Genomic Portal from Liver Hepatocellular Carcinoma (A) and Breast Cancer (B).

* indicates common proteins

A

| **liver tumor** | p Value |
|---|---|
| DKC1/NOP60b | 6.71E-15 |
| FBL | 0.000001171 |
| NOP10 | 0.00007354 |
| NOP16 | 0.000008451 |
| **NOP2*** | 1.59E-10 |
| NOP56 | 0.000001243 |
| NOP58 | 5.18E-14 |
| **RPS7*** | 5.54E-12 |
| RPS16 | 0.0008972 |
| RPS18 | 0.005751 |
| RPS20 | 0.0000206 |
| RPS21 | 0.0007177 |
| RPS27A | 1.23E-08 |
| RPS2P32 | 0.0000032 |
| RPSA | 0.004546 |
| RPL5 | 0.0005937 |
| RPL7 | 0.0191 |
| RPL21 | 0.0005989 |
| RPL24* | 0.00188 |
| RPL30 | 0.00006775 |
| **RPL35A*** | 0.0001787 |
| RPL38 | 0.0008094 |
| RPL39 | 0.00004626 |

B

| **brast tumor** | p Value |
|---|---|
| **NOP2*** | 0.0454 |
| **RPS7*** | 3.36E-04 |
| RPS8 | 0.0325 |
| RPL5 | 0.0444 |
| RPL12 | 0.0225 |
| RPL14 | 0.0389 |
| **RPL24*** | 0.0333 |
| RPL27 | 0.032 |
| RPL32 | 0.047 |
| RPL35 | 0.044 |
| **RPL35A*** | 0.0419 |

##### Fig. S7: List of PRIMERS used for qRT-PCRs

| gene | 5' FW sequence | 5' REV sequence | reference |
|---|---|---|---|
| *NOC1* | CTATACGCTCCACCGCACAT | GTCGCTACCGAACTTGTCCA | this work |
| *NOC2* | AGGAGCTTGAAGGGCTTAAAGA | ATCCTTGCTGGGTTTGTGGTA | this work |
| *NOC3* | TGCAGGCAGGCAAAAATCAC | AGCAAGCGTTTCATGAAGGC | this work |
| *E74b* | GAATCCGTAGCCTCCGACTGT | AGGAGGGAGAGTGGTGGTGTT | (Colombani et al., 2005) |
| *Actin5c* | CAGATCATGTTCGAGACCTTCAAC | ACGACCGGAGGCGTACAG | (Colombani et al., 2005) |
| *DILP8* | CGACAGAAG GTCCATCGAGT | GTT TTGCCG GATCCAAGTC | (Boulan et al., 2019) |
| *NOC1 genomic* | GTCACGGTCATTTCAATGGTA | CATGTCCAGCACCTCATC | this work |
| *ITS1* | GAAGAAACAAAATTCGAAAG | CGTATGCCCATAACTAAGAT | (Neumuller et al., 2013) |
| *ITS2* | ATCTTAGTTATGGGCATACG | CTGGCATATATCAATTCCTT | (Neumuller et al., 2013) |
| *18S* | CTCATATCCGAGGCCCTGTA | ACGAACGTTTTAACCGCAAC | (Neumuller et al., 2013) |
| *28S* | CGCTACGTCCGTTGGATTAT | CAATGCAAATTGCCCCTTAT | (Neumuller et al., 2013) |
| *XRP1* | GACCACACCGGAGATTATCAA | GCTGGTACTGGTACTTGTGGTG | (Baillon et al., 2018) |

**Baillon, L., Germani, F., Rockel, C., Hilchenbach, J. and Basler, K.** (2018). Xrp1 is a transcription factor required for cell competition-driven elimination of loser cells. *Scientific reports* **8**, 17712.

**Boulan, L., Andersen, D., Colombani, J., Boone, E. and Leopold, P.** (2019). Inter-Organ Growth Coordination Is Mediated by the Xrp1-Dilp8 Axis in Drosophila. *Dev Cell* **49**, 811-818 e814.

**Colombani, J., Bianchini, L., Layalle, S., Pondeville, E., Dauphin-Villemant, C., Antoniewski, C., Carre, C., Noselli, S. and Leopold, P.** (2005). Antagonistic actions of ecdysone and insulins determine final size in Drosophila. *Science* **310**, 667-670.

**Neumuller, R. A., Gross, T., Samsonova, A. A., Vinayagam, A., Buckner, M., Founk, K., Hu, Y., Sharifpoor, S., Rosebrock, A. P., Andrews, B., et al.** (2013). Conserved regulators of nucleolar size revealed by global phenotypic analyses. *Sci Signal* **6**, ra70.
